# A host ATPase essential for rhinovirus replication is an antiviral target with a high barrier to resistance

**DOI:** 10.64898/2026.05.13.723454

**Authors:** Matthew T. James, Courtney Dane, Karolina Wojtania, Caoimhe McAuley, Andrea Goya Grocin, Remigiusz A. Serwa, Michael Glenn, Erin Getty, Aidan O’Riain, Jack William Houghton, Aidan Ferris, Sheerien Manzoor, David G. Courtney, Ultan F. Power, Edward W. Tate, Aurelie Mousnier

## Abstract

Rhinoviruses are the leading cause of acute respiratory illnesses and comprise more than 170 types that constantly circulate in humans worldwide. Beyond common colds, rhinoviruses can trigger severe symptoms, particularly in young children, older adults and people with asthma or chronic obstructive pulmonary disease. Despite their clinical and socio-economic impact, no approved vaccine or antiviral treatment exist. Here, we uncovered the interaction of the host AAA+ ATPase RUVBL1/2 with rhinovirus non-structural protein 2C and we demonstrated that RUVBL1/2 is strictly and specifically required for the replication of the viral RNA of the most prevalent and pathogenic rhinovirus species. Pharmacological inhibition of RUVBL1/2 ATPase activity efficiently inhibited rhinovirus replication in a human nasal epithelium model, even post-infection. Moreover, serial viral passaging in the presence of a RUVBL1/2 inhibitor did not lead to the emergence of resistance. These findings reveal an unexpected and strong host dependency with promising potential for antiviral targeting.

## Introduction

Rhinoviruses (RVs) infect human populations worldwide throughout the year and are the leading cause of acute respiratory illnesses, often detected far more frequently than other respiratory viruses, even during the SARS-CoV-2 pandemic.^1–8^

Although RV infections are typically self-limiting and confined to the upper respiratory tract, their sheer frequency — often several episodes per year, particularly in children — imposes major health and socio-economic burdens. These stem from cumulative lost days of work/education, medical visits and antibiotic use, often without clearly diagnosed bacterial infection, potentially contributing to antibiotic resistance. Notably, a US cohort study reported that 23.5% of infected participants missed at least 1 day of work or school, 22.5% sought medical care, and 10.5% used antibiotics.^2^

Beyond causing common colds, which can lead to rhinosinusitis or otitis media, RVs are also responsible for a large portion of lower respiratory tract infections, which can require hospitalisation and may lead to death. RVs are indeed often the sole respiratory pathogens detected in such infections, and many studies have reported that RVs are among the most frequently identified respiratory viruses in hospitalised adults and children with respiratory infections.^9,10^ Notably, cohort studies have shown that RV-infected hospitalised older adults had a 30-day mortality rate of 8-10%.^11,12^ In addition, RVs are the dominant trigger for asthma exacerbations, detected in 51-71% of cases.^13^ These exacerbations frequently result in emergency department attendance and hospitalisation, especially for children, and can be fatal.^14^ RVs are also among the most common pathogens identified during acute exacerbations of chronic obstructive pulmonary disease (COPD), a frequent cause of hospitalisation and a key driver of morbidity and mortality among COPD patients.^9,15^

Despite their considerable clinical and socio-economic impact, there are currently no licensed vaccines or antiviral treatments for RV infections. Progress has been hampered by the extraordinary genetic diversity of RVs: over 170 (geno)types across 3 species (commonly named RV-A/B/C) that constantly co-circulate.^16^ Surveillance from Slovenia illustrates this, detecting 32–44% of all known RV types annually and most RV-A (91%) and RV-C (82%) types over an 8-year period.^1^ Although efforts to identify priority targets for vaccine or antiviral development have not yielded a clear shortlist of the most prevalent and clinically relevant types, they consistently show that RV-A and RV-C types dominate circulation and are associated with more severe disease than RV-B.^1,8,16,17^

Despite their diversity, most RVs rely on a common set of host proteins to replicate. Targeting these shared, host-dependent replication mechanisms offers a promising strategy for broadlZlspectrum antiviral development, unlikely to be easily affected by viral genetic variation. Several essential host factors have already been identified for RVs and related enteroviruses, such as poliovirus.^18–21^ One notable example is PI4KIIIβ,^22^ against which an inhibitor has recently entered a Phase 1 clinical trial (NCT05398198) in mild asthmatics experimentally infected with RV. Although results are not yet published, this underscores the potential of host proteins as antiviral targets.

Because our knowledge of the range of host proteins that support RV replication is likely incomplete, identifying additional host factors could improve our understanding of how RVs use host cells to replicate and broaden opportunities for antiviral development. We reasoned that such proteins could be uncovered through their interaction with components of viral replication complexes, where the viral RNA (vRNA) is replicated on modified cellular membranes through the coordinated activities of viral and cellular proteins. The viral non-structural protein 2C is an essential component of these complexes and plays a critical role in vRNA replication.^23^ Here, we analyse the 2C protein interactome in infected cells, revealing that host proteins RUVBL1 and RUVBL2 associate with 2C. We uncover an unexpected and essential role for these proteins in RV replication and show that an established RUVBL1/2 inhibitor prevents viral replication in infected cells, demonstrating the potential to target this mechanism pharmacologically.

## Results

### RUVBL1 and RUVBL2 interact with RV non-structural protein 2C and are required for viral replication

To identify host proteins that associate with RV 2C during infection, we infected HeLa-H1 cells with RV-A16 (MOI 20), formaldehyde cross-linked protein-protein interactions formed at 4.5 or 6 hours post-infection (hpi), immunoprecipitated the 2C-containing complexes, and analysed their composition by quantitative proteomics analysis (Figure 1A). Uninfected samples or samples immunoprecipitated with a non-2C-targeting antiserum served as controls. Several host proteins known to be recruited to the viral replication complexes or to play a role in enterovirus replication^19,22,24–26^ were significantly enriched in the 2C-containing fractions (e.g., PI4KIIIβ, SETD3, Arf1, Arf4, Arf5, GBF1), indicating isolation of genuine replication complexes (Figure 1B and 1C). Consistent with previous findings,^19,22^ knockdown of PI4KIIIβ and SETD3 with siRNA significantly inhibited RV-A16 replication (Figure 1D) without affecting cell viability (Figure 1E). Strikingly, although the RuvB-like proteins RUVBL1 and RUVBL2 have never been reported to be involved in the replication of any enterovirus, they were strongly enriched in the 2C-immunoprecipitated fractions as early as 4.5 hpi (Figure 1C), and their knockdown led to a nearly complete inhibition of RV-A16 replication (Figure 1D) without affecting cell viability (Figure 1E). Remarkably, knockdown of RUVBL1 or RUVBL2 had the most dramatic effect on RV-A16 replication, among all proteins enriched in the 2C-associated fractions with no significant impact on cell viability (Figure 1D and 1E).

**Figure 1.**
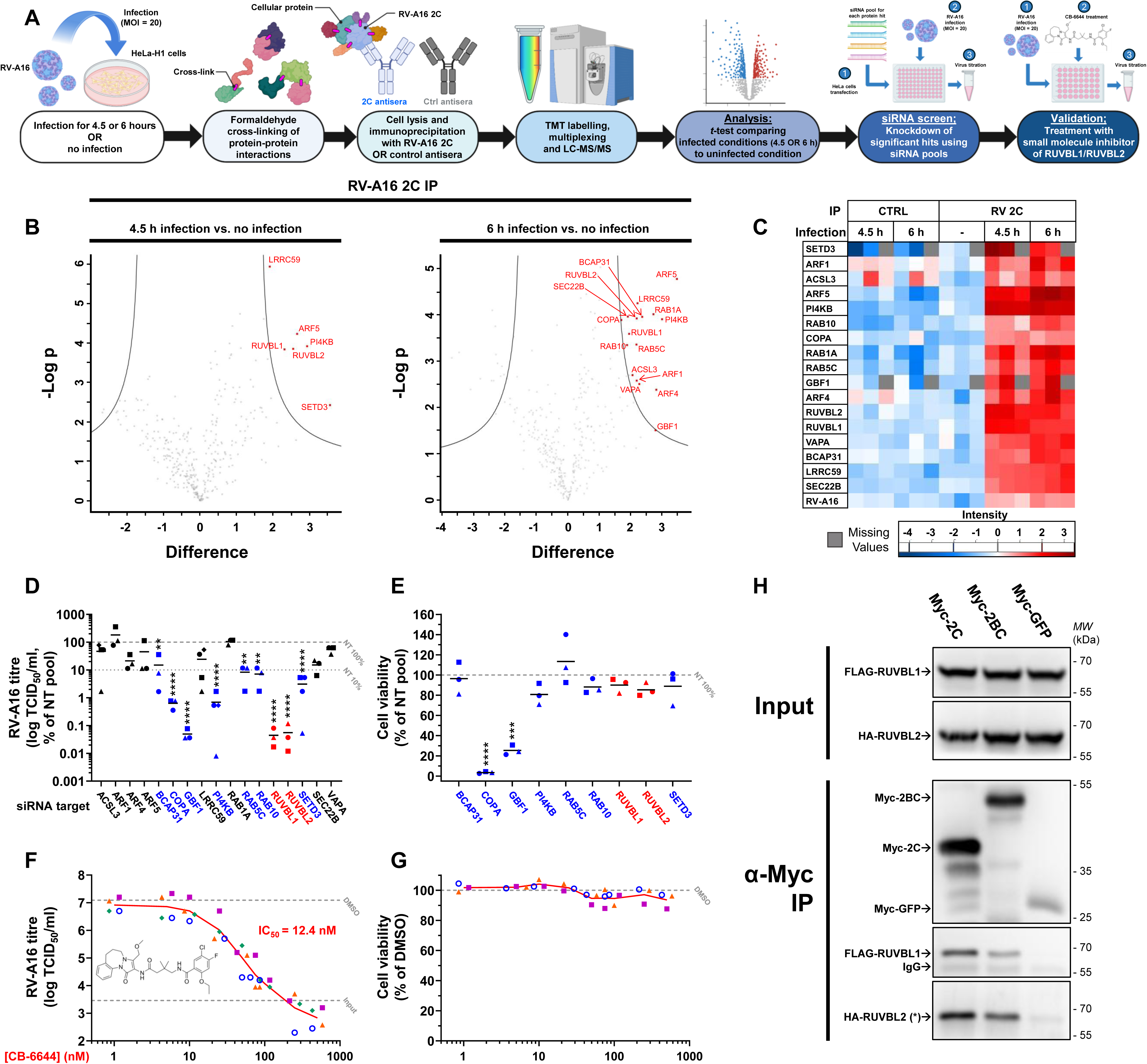
Analysis of rhinovirus 2C-associated proteins in infected cells reveals RUVBL1 and RUVBL2 as new cellular interactors critical for viral replication. **(A) Workflow for the identification of RV-A16 2C interactors and their functional validation.** HeLa-H1 cells were infected in triplicate with RV-A16 (MOI 20) for 4.5 h, 6 h or were left uninfected. Protein-protein interactions were then cross-linked with formaldehyde and cell lysates were immunoprecipitated using RV-A16 2C-specific or control antisera (2C-IP or control-IP). Eluates were TMT-labelled, multiplexed and analysed by LC-MS/MS. Differences between infected and uninfected conditions were assessed using two-sample Student’s *t*-test. Cellular proteins significantly enriched in infected 2C-IP fractions were further investigated for their role in RV-A16 replication through siRNA screening. The two most significant non-cytotoxic hits from the siRNA screen, RUVBL1 and RUVBL2, were further validated using a small molecule inhibitor. **(B-C) Proteomics analysis of 2C-IPs.** RV-A16 2C was immunoprecipitated from infected cells and the associated proteins were analysed as described in A. **(B)** Volcano plots showing in red the cellular proteins significantly enriched in 2C-IPs at 4.5 hpi (top) and 6 hpi (bottom), compared to uninfected conditions. **(C)** Corresponding heatmap, also showing infected control-IPs. **(D-E) siRNA screen of the 2C-IP hits.** HeLa-H1 cells were transfected with siRNA pools targeting the 2C-IPs hits or a non-targeting (NT) control siRNA pool. **(D)** At 72 h post-transfection, cells were infected with RV-A16 (MOI 20), and viral titres were quantified at 6 hpi. Data are presented as a percentage of the mean NT control (dashed grey line). **(E)** Viability of uninfected cells at 72 h post-transfection, for siRNA significantly reducing viral replication, presented as percentage of the mean NT control (dashed grey line). Statistical tests: one-way ANOVA with Dunnett’s post-hoc test, comparing to the NT control pool. **, *P* < 0.01; ***, *P* < 0.001; ****, *P* < 0.0001. **(F-G) Validation using CB-6644, a small molecule inhibitor of the ATPase activity of the RUVBL1/2 complex. (F)** HeLa-H1 cells were infected with RV-A16 (MOI 20) and treated at 1 hpi with DMSO or increasing concentrations of CB-6644. Viral titres were quantified at 0 hpi and 6 hpi. Viral titres in CB-6644-treated cells at 6 hpi are shown as individual points with means connected by a line. Mean viral titres of untreated cells at 0 hpi (input) and of DMSO-treated cells at 6 hpi are represented by dashed lines. **(G)** Cell viability of uninfected CB-6644-treated cells measured in parallel of the infection, presented as a percentage of the DMSO control. **(H) RUVBL1 and RUVBL2 co-immunoprecipitate with RV-A16 2C or 2BC in the absence of other viral components.** HeLa-H1 cells were transfected with constructs encoding FLAG-RUVBL1, HA-RUVBL2, and Myc-tagged RV NSPs (2C or 2BC) or Myc-GFP. Myc-tagged proteins were immunoprecipitated from cell lysates. Cell lysates (input) and immunoprecipitated fractions (α-Myc IP) were analysed by western blotting for Myc, FLAG, and HA. (*) HA-RUVBL2 overlaps with IgG heavy chain. FLAG-RUVBL1 and HA-RUVBL2 with Myc-2C, N=4; FLAG-RUVBL1 with Myc-2BC, N=3; HA-RUVBL2 with Myc-2BC, N=2. For all graph panels (D-G), data from 3-4 independent experiments are shown as individual points coded by shape, according to experimental replicate, together with means (connected by lines in F-G). Non-graph panels (H) show representative images. See also Figure S1.

RUVBL1 and RUVBL2 are related proteins of the AAA+ superfamily (ATPases associated with diverse cellular activities) that assemble into a hetero-hexameric complex, the RUVBL1-RUVBL2 complex (called RUVBL1/2 hereafter).^27^ RUVBL1/2 ATPase activity is blocked selectively by CB-6644, a small molecule allosteric inhibitor that interacts with ATP-bound RUVBL1/2 at the interface between two subunits, trapping it in a conformation that prevents ATP hydrolysis.^28,29^ Remarkably, when cells were treated with CB-6644, RV-A16 replication was potently inhibited, in a dose-dependent manner, with complete inhibition at 250-500 nM, a 50% inhibitory concentration (IC_50_) of 12.4 nM (95% confidence interval: 8.5 - 18.3 nM) (Figure 1F), and no significant effect on cell viability (Figure 1G). In agreement with the siRNA data (Figure 1D), these results demonstrate that the RUVBL1/2 ATPase activity is strictly required for RV-A16 replication in HeLa cells and can be targeted with a small molecule inhibitor.

Since RUVBL1 and RUVBL2 were identified as components of 2C-associated multi-protein complexes isolated from infected cells (Figure 1B and 1C), their interaction with 2C may have been mediated by other viral proteins and/or the vRNA. To determine if RUVBL1/2 could interact with 2C, or its uncleaved precursor 2BC, in the absence of other viral components, we evaluated their interaction in uninfected cells co-expressing the tagged proteins after transfection. In these conditions, RUVBL1 and RUVBL2 co-immunoprecipitated with RV-A16 2C or 2BC (Figure 1H), indicating that RUVBL1 and RUVBL2 can interact with 2C and 2BC independently of any other viral component.

Together, these results indicate that RUVBL1/2 interacts with RV-A16 2C/2BC and plays an essential role in the replication of the virus.

### CB-6644, a RUVBL1/2 inhibitor, prevents the replication of RV-A and RV-C types, in cell lines and in a human nasal epithelium model

To determine if RUVBL1/2 is a host factor required for the replication of other RV types beyond RV-A16, we tested the replication of other RV-A types (RV-A1b, RV-A29), which attach to host cells by binding to low-density lipoprotein receptor (LDLR), rather than intercellular adhesion molecule 1 (ICAM-1) required for RV-A16 entry. We also tested the replication of RV-B and RV-C types (RV-B14, RV-C15 and RV-C41). CB-6644 treatment completely inhibited the replication of all RV-A (Figure 2A, I-III) and RV-C (Figure 2C, I-II) types tested but did not affect RV-B14 replication (Figure 2A, IV), indicating that RUVBL1/2 is required for the replication of RV-A and RV-C, but not RV-B types. This result was not only observed in HeLa cells, but also in the BEAS-2B immortalised bronchial epithelial cell line, in which CB-6644 treatment inhibited the replication of RV-A types (Figure 2B, I-III) but not RV-B14 (Figure 2B, IV).

**Figure 2.**
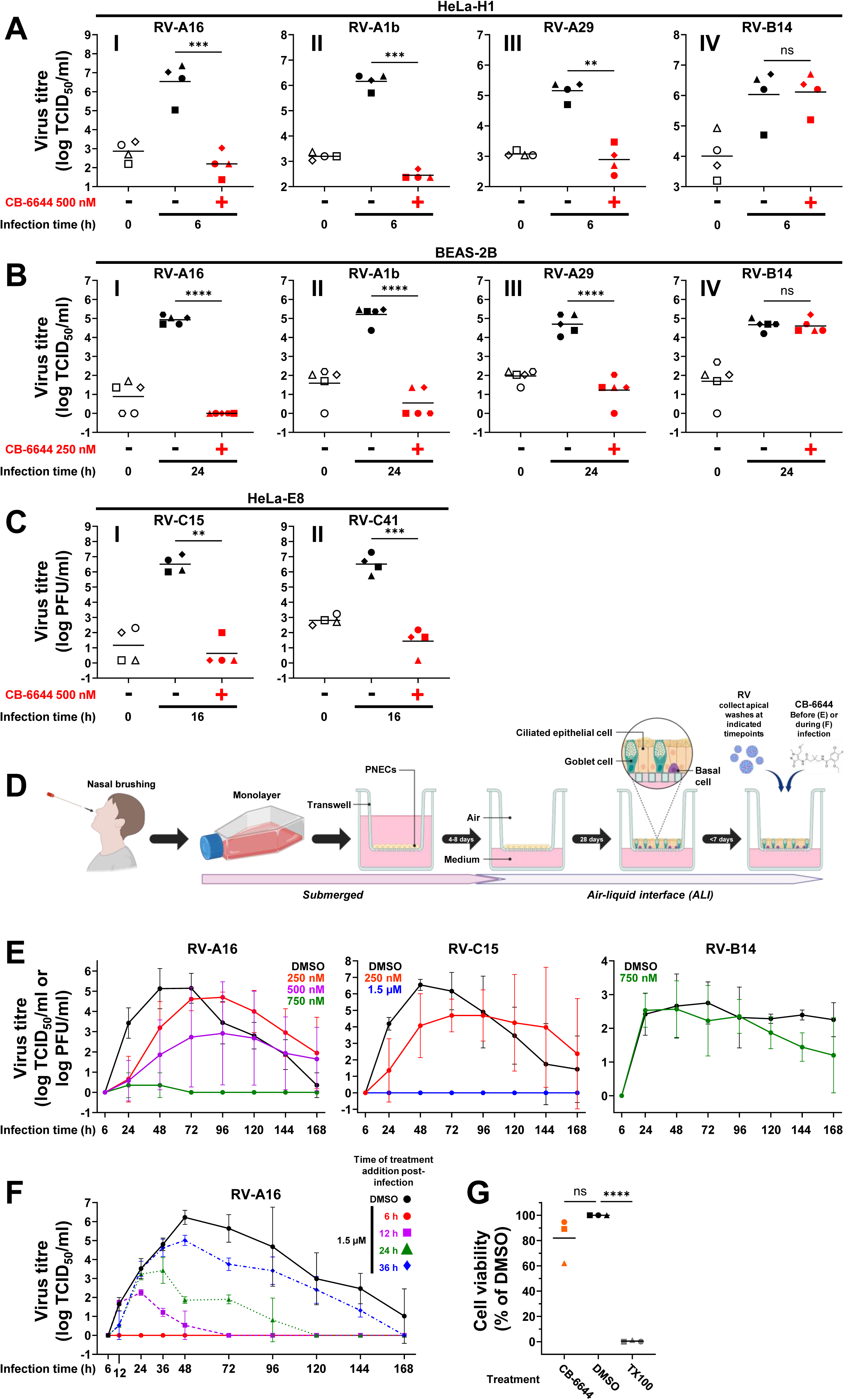
CB-6644 potently inhibits the replication of RV-A and RV-C types in cell lines and in well-differentiated primary nasal epithelial cell (WD-PNEC) cultures. **(A-C) CB-6644 antiviral assays in cell lines. (A)** HeLa-H1 cells, **(B)** BEAS-2B cells or **(C)** HeLa-E8 cells were infected with the indicated RV types (MOI 20 for **A**, MOI 1 for **B-C**). Cells were treated at 1 hpi with DMSO or the indicated concentrations of CB-6644. Viral titres were quantified at the indicated times post-infection. N=4 or 5 independent experiments. **(D-G) CB-6644 antiviral assays in WD-PNECs. (D)** Workflow for generation of WD-PNEC cultures. Primary nasal epithelial cells (PNECs) were sampled via nasal brushing from volunteers, expanded in monolayers, and seeded into Transwells. When 100% confluent, after 4-8 days of incubation, apical medium was removed to initiate air-liquid interface (ALI), which triggers cell differentiation and the formation of a pseudostratified epithelium containing ciliated epithelial cells, goblet cells and basal cells. After 28 days of incubation, high quality WD-PNEC cultures were infected apically with the indicated RV (MOI 0.01). CB-6644 or DMSO was added apically 16 h before **(E)** or at different time points after **(F)** infection, as indicated. Viral titres in apical washes collected at the indicated times were quantified. N= 3 (E, RV-A16 and RV-B14) or 2 (E, RV-C15 and F) independent donors. **(G)** Viability of WD-PNECs apically treated with 2 μM CB-6644 or DMSO for 192 h, or with 1% Triton X-100 (TX100) for 2 h, presented as percentage viability relative to DMSO-treated control. N= 3 independent donors. For panels A-C and G, data are shown as individual points, coded by shape according to experimental replicate, with means. For panels E-F, data are shown as means (± SD) connected by lines colour-coded by treatment. Statistical tests: two-tailed paired *t*-test (A-C), one-way ANOVA with Dunnett’s *post-hoc* test (G). **, *P* < 0.01; ***, *P* < 0.001; ****, *P* < 0.0001; ns, not significant. See also Figure S2.

Similar results were observed in well-differentiated primary human nasal epithelial cell cultures (WD-PNECs) derived from nasal brushes from healthy adult volunteers (Figure 2D-G). WD-PNECs are physiologically and morphologically authentic relative to the nasal epithelium *in vivo*.^30,31^ Consistent with a time-course of RV release in nasal secretions of infected volunteers,^32,33^ and in bronchial epithelium models,^34,35^ peak viral titres were detected between 48 and 72 hpi in DMSO-treated control WD-PNECs (Figure 2E). Pre-treatment of WD-PNECs with CB-6644 inhibited RV-A16 and RV-C15 replication, with complete inhibition at the highest concentrations (750 nM for RV-A16 and 1.5 µM for RV-C15, Figure 2E). In contrast, as observed in HeLa and BEAS-2B cell lines, RV-B14 replication in WD-PNECs was not inhibited by CB-6644 (Figure 2E). Interestingly, CB-6644 inhibited RV-A16 replication even when added after the start of the infection (Figure 2F), demonstrating significant therapeutic potential even when viral titres are already high. Indeed, viral titres diminished rapidly upon CB-6644 addition, indicating efficient inhibition of further rounds of replication (Figure 2F). Cell viability in WD-PNEC cultures was not affected by CB-6644, even after 192 hours treatment with 2 µM CB-6644 (Figure 2G).

Together, these results indicate that RUVBL1/2 activity is strictly required for the replication of RV-A and RV-C types, but not RV-B types, in both cell lines and in an authentic model of the human nasal epithelium.

### RUVBL1/2 is required for synthesis of RV negative-strand RNA, after translation of the incoming viral positive-strand RNA

To investigate the mechanism by which RUVBL1/2 plays an essential role in the RV replication cycle, we first examined the effect of its inhibition or knockdown on the synthesis of vRNA and viral proteins. Treatment of HeLa cells with CB-6644 or siRNA against RUVBL1, which knocks down expression of both RUVBL1 and RUVBL2 (Figure S2B-C) as previously reported,^29^ inhibited the synthesis of RV-A16 vRNA (Figure 3A and 3E) and 3C and 2C non-structural proteins (Figure 3B-C, 3F-J, S2D-F). This indicates that RUVBL1/2 is required at an early stage of the viral replication cycle. To determine whether RUVBL1/2 was required after viral entry, we assessed the effect of CB-6644 treatment when normal viral entry was bypassed by direct transfection of the vRNA into cells. Following vRNA transfection, RV-A16 and RV-A1a virion production was totally inhibited by CB-6644 treatment (Figure 3K), demonstrating that RUVBL1/2 was indispensable for RV-A replication after entry. In contrast, and consistent with previous infection experiments (Figure 2A-B, IV), CB-6644 did not inhibit RV-B14 virion production following vRNA transfection (Figure 3K). CB-6644 time of addition experiments support a role for RUVBL1/2 in the early stages of the replication cycle after entry, as CB-6644 fully inhibited viral replication when added at 1 hpi, but the inhibitory effect was progressively lost when added later (Figure 3D).

**Figure 3.**
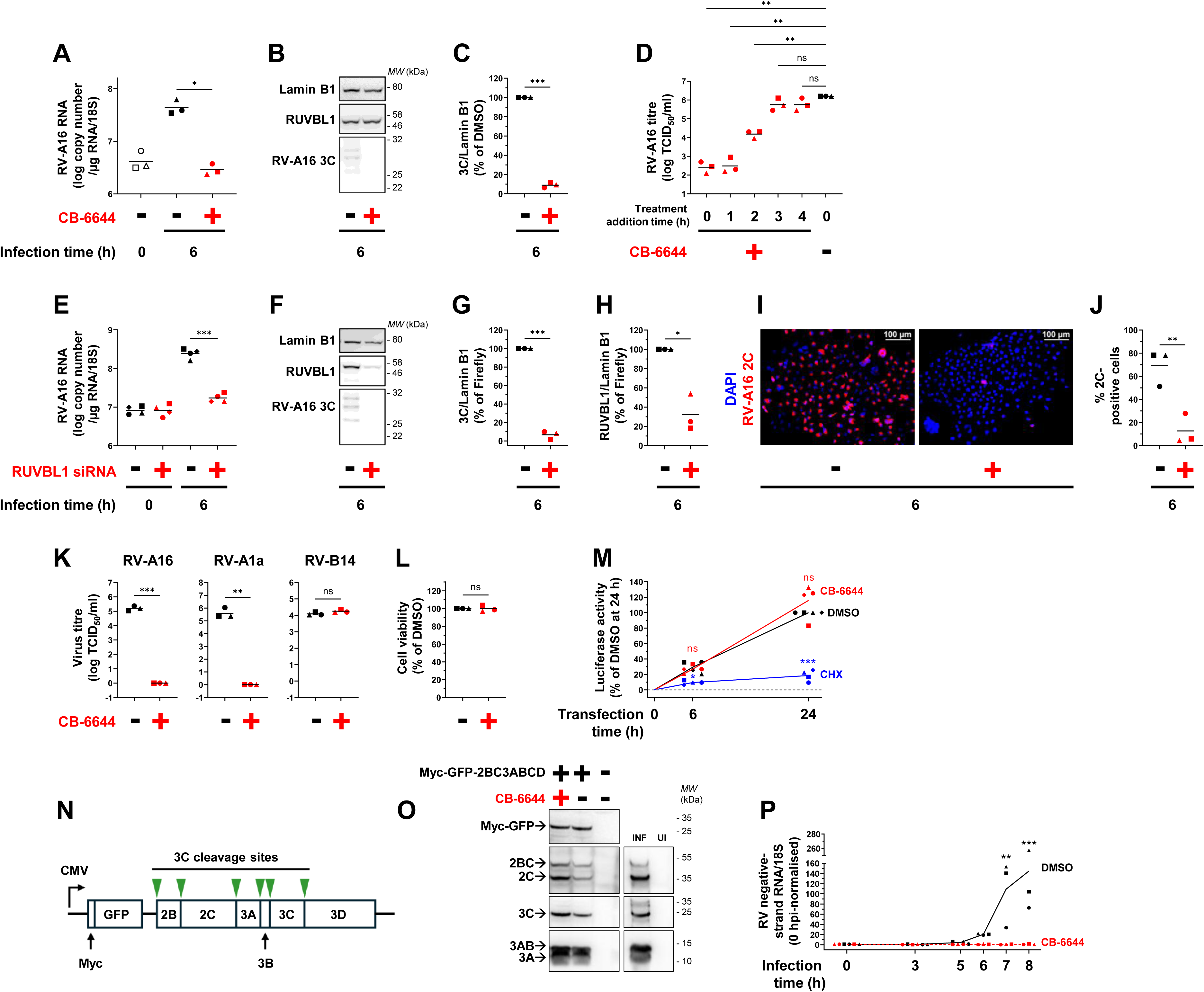
RUVBL1/2 is required after viral entry for RV RNA replication, but not IRES-dependent translation or polyprotein processing. **(A–C) CB-6644 inhibits RV RNA replication and NSP production.** HeLa-H1 cells were infected with RV-A16 (MOI 20) and treated at 1 hpi with DMSO or 500 nM CB-6644. **(A)** Viral RNA was quantified by RT-qPCR at 0 hpi and 6 hpi. **(B)** At 6 hpi, lysates were analysed by western blotting for RV-A16 3C, RUVBL1, and lamin-B1. **(C)** 3C signal was quantified and normalised to lamin-B1. **(D) Time-of-addition assay.** HeLa-H1 cells were infected as above and treated with DMSO or 500 nM CB-6644 immediately after virus adsorption (0 hpi) or at the indicated times post-infection. Viral titres were quantified at 6 hpi. **(E–J) siRNA knockdown of RUVBL1 inhibits RV RNA replication and NSP production.** HeLa-H1 cells were transfected with siRNA targeting RUVBL1 or firefly luciferase for 72 h and then infected with RV-A16 (MOI 20). **(E)** Viral RNA was quantified by RT-qPCR at 0 hpi and 6 hpi. **(F)** At 6 hpi, lysates were analysed by western blotting for RV-A16 3C, RUVBL1, and lamin-B1. **(G–H)** Quantification of 3C and RUVBL1 signal from F, normalised to lamin-B1. **(I)** Immunofluorescence staining for RV-A16 2C (red) at 6 hpi; nuclei were stained with DAPI (blue). **(J)** Quantification of 2C-positive cells from (I). **(K–L) RUVBL1/2 is required after RV entry. (K)** HeLa-H1 cells were transfected with RV-A16, RV-A1a, or RV-B14 RNA in the presence of DMSO or 500 nM CB-6644. Viral titres were quantified at 14 h post-transfection. **(L)** Cell viability assessed in parallel of K in untransfected cells treated for 14 h with DMSO or 500 nM CB-6644. **(M) RUVBL1/2 is not required for IRES-dependent translation.** HeLa-H1 cells were transfected with a luciferase reporter RNA under RV-A16 IRES-dependent translational control, in the presence of DMSO, 500 nM CB-6644, or cycloheximide (CHX). Luciferase activity was measured at the indicated times. Values were t=0-subtracted and normalised to the DMSO 24 h post-transfection value within each experiment. The 0 h baseline is shown as a dashed grey line. **(N-O) RUVBL1/2 is not required for RV-A16 polyprotein cleavage. (N)** Myc-GFP-2BC3ABCD construct used for polyprotein processing assays, with expression under the control of a CMV promoter. **(O)** HeLa-H1 cells were transfected or not with the Myc-GFP-2BC3ABCD plasmid for 21 h, in the presence of DMSO or 500 nM CB-6644. In parallel, HeLa-H1 cells were infected or not with RV-A16 for 8 h. Lysates were then analysed by western blotting for Myc-GFP and RV-A16 2C, 3A and 3C. **(P) CB-6644 inhibits negative-strand RNA synthesis.** HeLa-H1 cells were infected with RV-A16 (MOI 20) and treated with DMSO or 500 nM CB-6644 at 1 hpi. Negative-strand RNA was quantified at the indicated times by RT-qPCR, normalised to 0 hpi. For all graph panels (A, C-E, G, H, J-M, P), data from 3-4 independent experiments are shown as individual points, coded by shape according to experimental replicate, with means (connected by lines in M and P). Non-graph panels (B, F, I, O) show representative images from 3 independent experiments. Statistical tests: two-tailed paired t-test (A, C, E, G, H, J-L), one-way ANOVA with Dunnett’s *post-hoc* test (D), two-way ANOVA, comparing drug treatments to the DMSO control at each time point (M, P). *, *P* < 0.05; **, *P* < 0.01; ***, *P* < 0.001; ns, not significant. See also Figure S2 and S3.

Once released into the cell cytosol, the vRNA is directly translated by host ribosomes into a polyprotein, which is then cleaved by the viral proteases. Translation is mediated by an internal ribosome entry site (IRES), a structured RNA element present in the 5’ untranslated region (UTR) of the vRNA, which allows cap-independent translation. Following translation, the incoming vRNA is replicated and new copies are used for further translation. Viral proteins are only detectable by western blot or immunofluorescence from 4-5 hpi, once the vRNA available for translation has been sufficiently amplified through replication.^21^ Therefore, even though 3C and 2C non-structural protein are minimally detected when RUVBL1/2 is inhibited or knocked down (Figure 3B-C and 3F-J), this does not discriminate between direct inhibition of translation and an indirect consequence of the inhibition of vRNA replication (as observed in Figure 3A and 3E). To specifically assess the effect of CB-6644 on IRES-dependent translation, we transfected a luciferase reporter RNA that cannot self-replicate, in which luciferase expression was under the control of the RV-A16 IRES. While cycloheximide, a protein translation inhibitor, blocked the expression of the luciferase reporter, CB-6644 did not (Figure 3M), indicating that RUVBL1/2 inhibition does not prevent IRES-dependent translation.

Since RUVBL1/2 was required for vRNA replication (Figure 3A and 3E) but not translation (Figure 3M), we investigated whether it was necessary for cleavage of the viral polyprotein, a prerequisite for vRNA replication. Because viral proteins cannot be efficiently detected by western blot when infected cells are treated with CB-6644 (Figure 3B-C), even after immunoprecipitation (Figure S3B-E), we transfected cells with a plasmid allowing the expression of a RV-A16 polyprotein fragment containing the non-structural proteins 2B-2C-3A-3B-3C-3D, under ectopic transcriptional and translational control (CMV promoter, cap-dependent translation). A Myc-GFP reporter, which can be cleaved by the 3C protease, was placed upstream of the polyprotein fragment, to control for protein expression (Figure 3N). With this artificial expression system, cleavage of the polyprotein fragment into 2BC, 2C, 3AB, 3A and 3C was observed with or without CB-6644, similarly to untreated infected cells (Figure 3O). This indicates that RUVBL1/2 inhibition does not prevent polyprotein cleavage, and it is corroborated by the detection of cleaved 3C, 3A/3AB and 2C after immunoprecipitation of the small amount of viral proteins synthesised in infected cells treated with CB-6644 (Figure S3B-D). Together, these results indicate that RUVBL1/2 is not essential for polyprotein cleavage, suggesting that it may be directly involved in vRNA replication.

A standard RT-qPCR assay, which mostly detects the positive-strand vRNA genome, demonstrated that CB-6644 inhibits RV-A16 vRNA replication (Figure 3A and 3E). To further assess if CB-6644 interfered with the first step of vRNA replication, the synthesis of negative-strand copies of the viral genome which are then used as templates for positive-strand RNA synthesis, we used an RT-qPCR assay that selectively quantifies negative-strand RV-A16 RNA.^35,36^ In DMSO control conditions, negative-strand vRNA synthesis was clearly detected between 6 and 8 hpi, whereas it was completely abrogated by CB-6644 treatment (Figure 3P).

Taken together, these data indicate that following RV-A16 entry and translation of the incoming vRNA, RUVBL1/2 plays an essential role in negative-strand vRNA synthesis, but it is not required for viral polyprotein cleavage.

### CB-6644 inhibits RV replication independently of cellular transcription and resistance does not readily emerge

Since CB-6644 treatment upregulates the transcription of hundreds of genes, including interferon response genes,^29,37^ the possibility remained that the antiviral activity of CB-6644 was indirect, for example through upregulation of interferon stimulated genes (ISGs). However, inhibition of transcription (with actinomycin D or triptolide) or of the JAK-STAT pathway (with ruxolitinib, which blocks interferon-induced ISG transcription) did not restore vRNA replication in cells treated with CB-6644 (Figure 4A). In contrast, treatment of cells with actinomycin D, triptolide or ruxolitinib, following stimulation of the interferon response with poly(I:C), effectively inhibited transcription of the ISG OAS1 (Figure S4A), demonstrating functional inhibition. These results indicate that the effect of CB-6644 on RV replication is not mediated through indirect effects involving the stimulation of interferon-mediated responses or the transcription of cellular genes. It is also not mediated by nonsense-mediated mRNA decay (NMD), in which RUVBL1/2 are involved,^38,39^ since the NMD inhibitors NMDI-14 and wortmannin did not restore viral replication in CB-6644-treated cells (Figure S4B-E). Together, these results are consistent with a direct role of RUVBL1/2 in vRNA replication.

**Figure 4.**
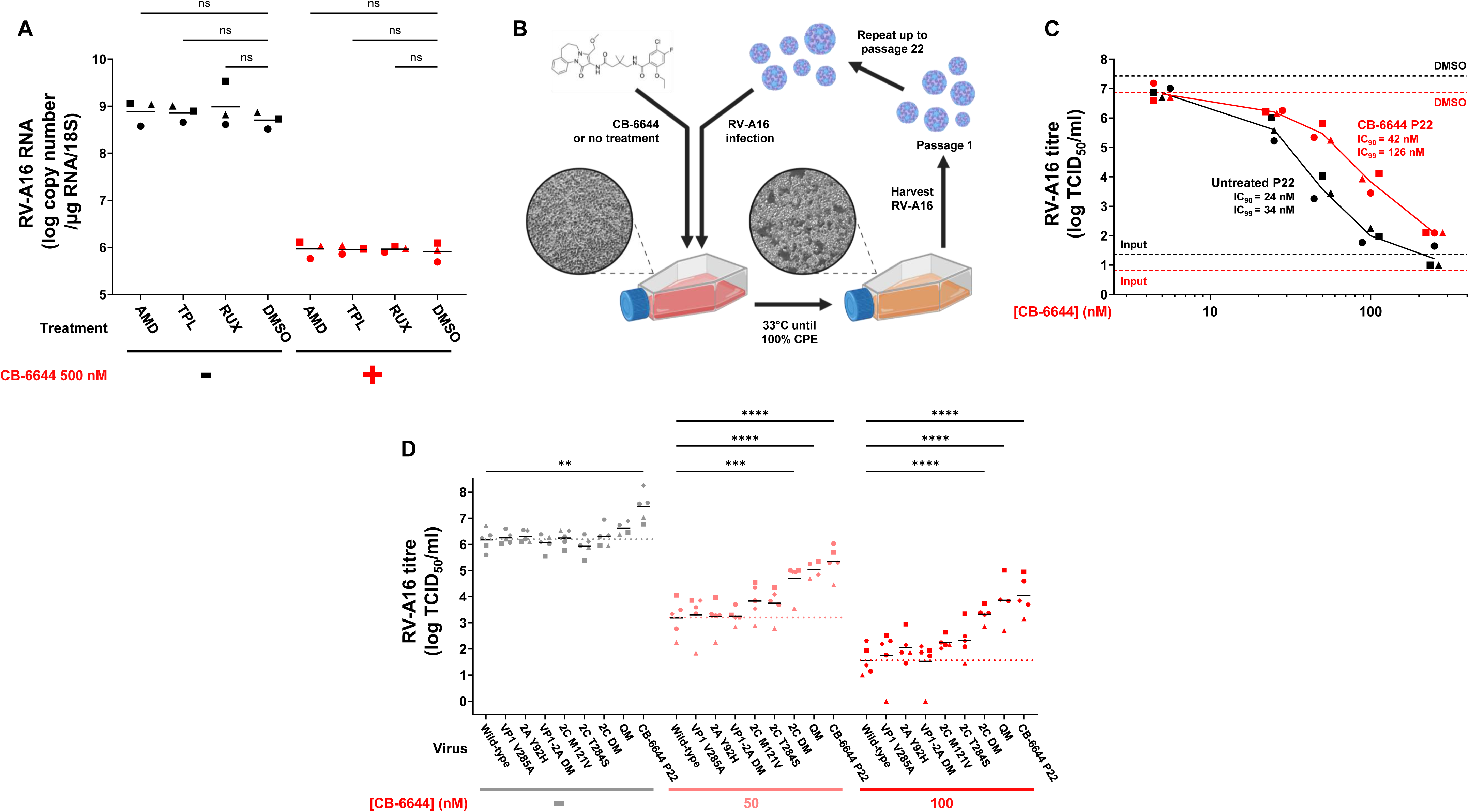
Mechanisms of RV-A16 sensitivity to CB-6644. **(A) Inhibitors of cellular transcription or of JAK1/2 do not abrogate the antiviral effect of CB-6644.** HeLa-H1 cells were pre-treated for 1 h with actinomycin D (AMD), triptolide (TPL), ruxolitinib (RUX), or DMSO, and subsequently infected with RV-A16 (MOI 20) in the presence of the corresponding drug and DMSO or 500 nM CB-6644. Viral RNA was quantified by RT-qPCR at 6 hpi (N=3). **(B) Generation of CB-6644-resistant RV-A16.** HeLa-H1 cells were infected with RV-A16 (MOI 0.1) in the presence of 15 nM CB-6644 or without treatment and incubated until 100% cytopathic effect was observed. Passage 1 virus was harvested and used to infect fresh HeLa-H1 cells under the same conditions. This process was repeated until passage 22 (P22), with CB-6644 concentrations being increased stepwise (30 nM at P3, 60 nM at P16 and 100 nM at P19). **(C) CB-6644-passaged RV-A16 exhibits reduced sensitivity to CB-6644.** HeLa-H1 cells were infected with passaged RV-A16 (P22, from untreated [black] or CB-6644-treated [red] passages, MOI 1) and were treated with DMSO or increasing concentrations of CB-6644. Viral titres were quantified at 0 hpi and 16 hpi (N=3). Viral titres in CB-6644-treated cells at 16 hpi are shown as individual points with means connected by lines. Mean viral titres of untreated cells at 0 hpi (input) and of DMSO-treated cells at 16 hpi are represented by dashed lines. **(D) Mutations in RV-A16 2C reduce sensitivity to CB-6644.** HeLa-H1 cells were infected with wild-type RV-A16, CB-6644-passaged virus (P22, from B), or with recombinant RV-A16 viruses carrying a single mutation (VP1 V285A, 2A Y92H, 2C M121V, or 2C T284S), two mutations (VP1 V285A + 2A Y92H [VP1-2A DM], or 2C M121V + T284S [2C DM]), or all four mutations in combination (quadruple mutant [QM]) (MOI 1). Infected cells were treated with DMSO or the indicated concentrations of CB-6644, and viral titres were quantified at 16 hpi (N=4). Dashed lines indicate mean titres of wild-type RV-A16 under each treatment. For all panels, N=number of independent experiments. For all graph panels (A, C-D), data are shown as individual points, coded by shape according to experimental replicate, with means (connected by lines in C). Statistical tests: one-way ANOVA with Dunnett’s *post-hoc* test (A), two-way ANOVA with Holm-Sidak’s *post-hoc* test (D). For D, only statistically significant comparisons are shown; all other comparisons within each treatment group were non-significant. **, *P* < 0.01; ***, *P* < 0.001; ****, *P* < 0.0001; ns, not significant. See also Figure S4.

To determine whether a RUVBL1/2-dependent RV type could evolve resistance to CB-6644, we serially passaged RV-A16 in the presence of increasing concentrations of CB-6644 (Figure 4B). After 22 passages, the virus only became slightly less sensitive to CB-6644 (Figure 4C). In contrast, after only 15 passages, RV-A16 developed significant resistance to the adenosine analogue NITD008 (Figure S4F), which inhibits enterovirus RNA synthesis through chain termination.^40,41^ Sequencing of the virus passaged in the presence of CB-6644 revealed mutations in VP1, 2A and 2C, compared to the parental virus (Figure 4D). While mutations in VP1 and 2A (alone or combined) did not alter sensitivity to CB-6644, the combined mutations in 2C (M121V and T284S) reduced sensitivity to CB-6644 to a level comparable to the passaged virus or the engineered quadruple mutant virus with mutations in VP1, 2A and 2C (Figure 4D). This indicates that RV-A16 strongly depends on the RUVBL1/2 AAA+ ATPase to replicate and that there is a high barrier to the development of resistance, likely requiring multiple mutations. The emergence of mutations in 2C (M121V in the β1-strand of the ATPase domain and T284S in the β6-strand of the zinc finger domain) during passaging with CB-6644, which slightly reduced RV sensitivity to the inhibitor, suggests selective pressure to mutate 2C to develop resistance.

## Discussion

Picornaviruses — the family of viruses that includes RVs — have among the smallest viral genomes (6.7–10.1 kb), encoding only ∼2,200 amino acids. Consequently, they depend even more than some other viruses on host cells to replicate, yet the host factors required and the underlying molecular mechanisms remain poorly defined. Elucidating these host dependencies can provide attractive targets to develop novel antivirals, with the concomitant advantage of reduced risks of resistance compared to direct-acting antivirals. Our study addresses this knowledge gap by uncovering previously uncharacterized host factors essential for rhinovirus replication and defining their role. This provides exciting new insights into picornavirus-host interactions and highlights exploitable vulnerabilities for therapeutic intervention.

By analysing RV-A16 2C-containing complexes isolated from infected cells, we identified known replication factors, which validated the isolation of authentic complexes. We also uncovered robust early recruitment of RUVBL1 and RUVBL2 to 2C-containing complexes. While other proteins not previously implicated in picornavirus replication (e.g., BCAP31) were also identified, knockdown of RUVBL1 or RUVBL2 had the most drastic effect on RV-A16 replication, nearly abolishing it. Furthermore, a selective inhibitor of RUVBL1/2 fully blocked viral replication in cell lines and in an authentic human nasal epithelium model. Given the striking impact of RUVBL1/2 knockdown or inhibition on RV-A and RV-C replication, it is surprising that RUVBL1 was only a low-ranked hit in previous siRNA and CRISPR-Cas9 screens for RV-A2 or RV-C15 proviral host factors.^19,42^ This likely reflects a limitation of genetic screens, where true proviral factors can be obscured by the large number of genes whose loss indirectly impairs viral replication. A similar issue may explain why the well-established enterovirus dependency factor PI4KIIIβ, which was strongly enriched in our 2C complexes, was ranked only 1108^th^ in a prior CRISPR-Cas9 screen.^19^ These observations highlight the strength of our direct interactomic approach to identify host factors genuinely engaged in viral replication complexes.

Mechanistically, our data indicate that following viral entry and translation of the incoming vRNA, RUVBL1/2 ATPase activity is essential for negative-strand vRNA synthesis, but dispensable for viral polyprotein cleavage. Co-immunoprecipitation experiments show that RUVBL1 and RUVBL2 associate with 2C/2BC in infected cells and in cells expressing 2C or 2BC without other viral components. In addition, virus passaging experiments in the presence of a RUVBL1/2 inhibitor suggest selective pressure to mutate 2C to develop resistance. Together, these data indicate that RUVBL1/2 engages with 2C/2BC to promote vRNA replication. In line with its role in the assembly and maturation of cellular protein and ribonucleoprotein complexes,^27,43^ RUVBL1/2 may participate in the assembly and remodeling of the viral replication complexes following vRNA translation to allow formation of specific protein complexes on the vRNA that enable initiation of negative-strand vRNA synthesis. This process indeed involves the coordinated recruitment of specific viral and cellular proteins to RNA structures located at the extremities and in the coding sequence of the vRNA, in association with cellular membranes.^44–49^ While the precise role of 2C in this process is not fully understood, studies on poliovirus have demonstrated that 2C binds to the vRNA and cellular membranes and plays an essential role in the initiation of negative-strand vRNA synthesis.^23,50–53^ Our results demonstrate that RUVBL1/2 interacts with RV-A16 2C, and its activity is required for negative-strand vRNA synthesis. We propose that, after translation of the vRNA, RUVBL1/2 cooperates with 2C to assemble membrane-associated ribonucleoprotein complexes that are competent for vRNA replication, potentially contributing to the switch from vRNA translation to replication by remodeling the viral ribonucleoprotein complexes.

Remarkably, although the RUVBL1/2 inhibitor CB-6644 completely blocked the replication of RV-A and RV-C types, it did not affect RV-B14 replication. This argues against broad cellular disturbances that would prevent any viral/enteroviral replication, rather pointing to a specific dependency. The shared reliance of RV-A and RV-C types on RUVBL1/2, contrasted with the independence of RV-B14, is consistent with established phylogenetic relationships in which RV-A and RV-C types are genetically more closely related to each other than to RV-B types.^54,55^ Interestingly, unlike RV-B14, RV-A and RV-C vRNA replication also depends on the host protein STING in a 2C-linked manner.^20^ Together, these findings suggest that, to support vRNA replication, RV-A and RV-C types engage with a distinct set of host proteins, compared to the more evolutionary distant RV-B types. We speculate that RV-B vRNA replication may require a different cellular AAA+ ATPase than RUVBL1/2, such as VCP/p97, which has been shown to be involved in poliovirus replication.^56,57^

We demonstrated the promising antiviral potential of targeting RUVBL1/2, using CB-6644, a small-molecule inhibitor of RUVBL1/2 originally developed for potential cancer treatment.^29^ CB-6644 potently and efficiently inhibited RV-A and RV-C replication in cell lines and in a human nasal epithelium model. Notably, no significantly resistant viruses emerged after 22 sequential passages in the presence of increasing concentrations of CB-6644. Although two mutations arose in 2C during passaging and conferred a modest reduction in sensitivity to CB-6644, the replication of the mutant virus remained strongly inhibited by CB-6644, with reductions in viral titres of around 3 log_10_ at 100 nM CB-6644, and 5 log_10_ at 250 nM CB-6644, compared with the DMSO-treated control. This indicates that the acquisition of resistance would likely require additional mutations. This conclusion is further supported by sequence comparisons of 2C from CB-6644-resistant RV-B14 and from CB-6644-sensitive RV-A16 and RV-C15. RV-B14 2C differs from RV-A16 and RV-C15 2C by 169 and 183 amino acid residues, respectively (109 and 110 non-similar residues). While not all these differences necessarily determine CB-6644 sensitivity, their extent, together with the passaging data, suggests that the development of resistance in RV-A and RV-C types would require the accumulation of many mutations, potentially linked to the acquisition of the ability to interact with different host cell proteins. Collectively, our results indicate that targeting RUVBL1/2 is a robust antiviral strategy with a high barrier to resistance.

Interestingly, although RUVBL1 and RUVBL2 have not been previously implicated in picornavirus replication, the knockdown of RUVBL2 has been shown to significantly reduce influenza A virus (IAV) RNA replication.^58^ This indicates that these AAA+ ATPase are not only important for RV-A and RV-C replication, but also for IAV, suggesting a broader role in the replication of several respiratory viruses. Consequently, CB-6644 may have antiviral activity against multiple clinically important respiratory viruses, underscoring its potential against viruses that pose a substantial clinical and public health burden.

In conclusion, we reveal an unexpected essential role for RUVBL1/2 in the replication of the most prevalent and pathogenic rhinovirus species. By mechanistically positioning RUVBL1/2 at the heart of RV-A/C vRNA replication and demonstrating the ability to pharmacologically inhibit this host-dependent process with a high resistance barrier, this work refines the picornaviral host-dependency map and opens new avenues for antiviral development.

## Resource availability

### Lead contact

Requests for further information and resources should be directed to and will be fulfilled by the lead contact, Aurelie Mousnier.

### Materials availability

All unique/stable reagents generated in this study are available from the lead contact with a completed materials transfer agreement.

### Data and code availability

The mass spectrometry proteomics data have been deposited to the ProteomeXchange Consortium via the PRIDE^59^ partner repository with the dataset identifier PXD075813.

## Acknowledgments

We thank Sebastian Johnston and Roberto Solari for the gift of plasmid pCR2.1_RV-A16 and antibodies against RV-A16 non-structural proteins 2C, 3A, 3C and 3D. We thank James Gern and Yury Bochkov for the gift of HeLa-E8 cells and plasmids pC15-Rz-K41 and pC41-Rz-K41. We thank Stanley Lemon and Kevin McKnight for the gift of plasmids pR16.11, pWIN1a, pWR3.26 and pRVA-16-NL. We thank Gad Frankel and Gunnar Schroeder for the gift of plasmid pICC1564. We thank Steven Artandi for the gift of plasmids pCDNA-3xFLAG-Pontin and pcDNA-3xHA-Reptin.

This work was supported by the Medical Research Foundation and Asthma + Lung UK [grant number MRFAUK-2015-311]; the Medical Research Council [grant number MR/X020371/1]; and a Medical Research Council Confidence in Concept grant [grant number CD1920 - CIC12]. In addition, M.T.J. and C.D. were each supported by a studentship from the Department for the Economy of the Northern Ireland Executive, and D.G.C. was supported by an ERC-StG grant [grant number PTFLU 949506].

## Author contributions

Conceptualization, A.M. and M.T.J.; Data Curation, M.T.J., C.D., A.G.G., R.A.S., M.G., J.W.H., A.F., D.G.C., A.M.; Formal Analysis, M.T.J., C.D., A.G.G., R.A.S., M.G., J.W.H., A.F., D.G.C., A.M.; Funding Acquisition, A.M.; Investigation, M.T.J., C.D., K.W., C.M., A.G.G., R.A.S., M.G., E.G., A.O., A.F., D.G.C.; Methodology, M.T.J., C.D., A.G.G., R.A.S., M.G., S.M., D.G.C., U.F.P., E.W.T., A.M.; Project Administration, A.M., E.W.T., U.F.P.; Resources, E.W.T., U.F.P., M.G., D.G.C.; Supervision, A.M., E.W.T., U.F.P.; Visualization, M.T.J., A.G.G., J.W.H.; Writing – original draft, A.M. and M.T.J.; Writing – review & editing, C.D., M.G., J.W.H., A.F., D.G.C, U.F.P., E.W.T.

## Declaration of interests

The authors declare no competing interests.

## STAR Methods

### 1 Key resources table

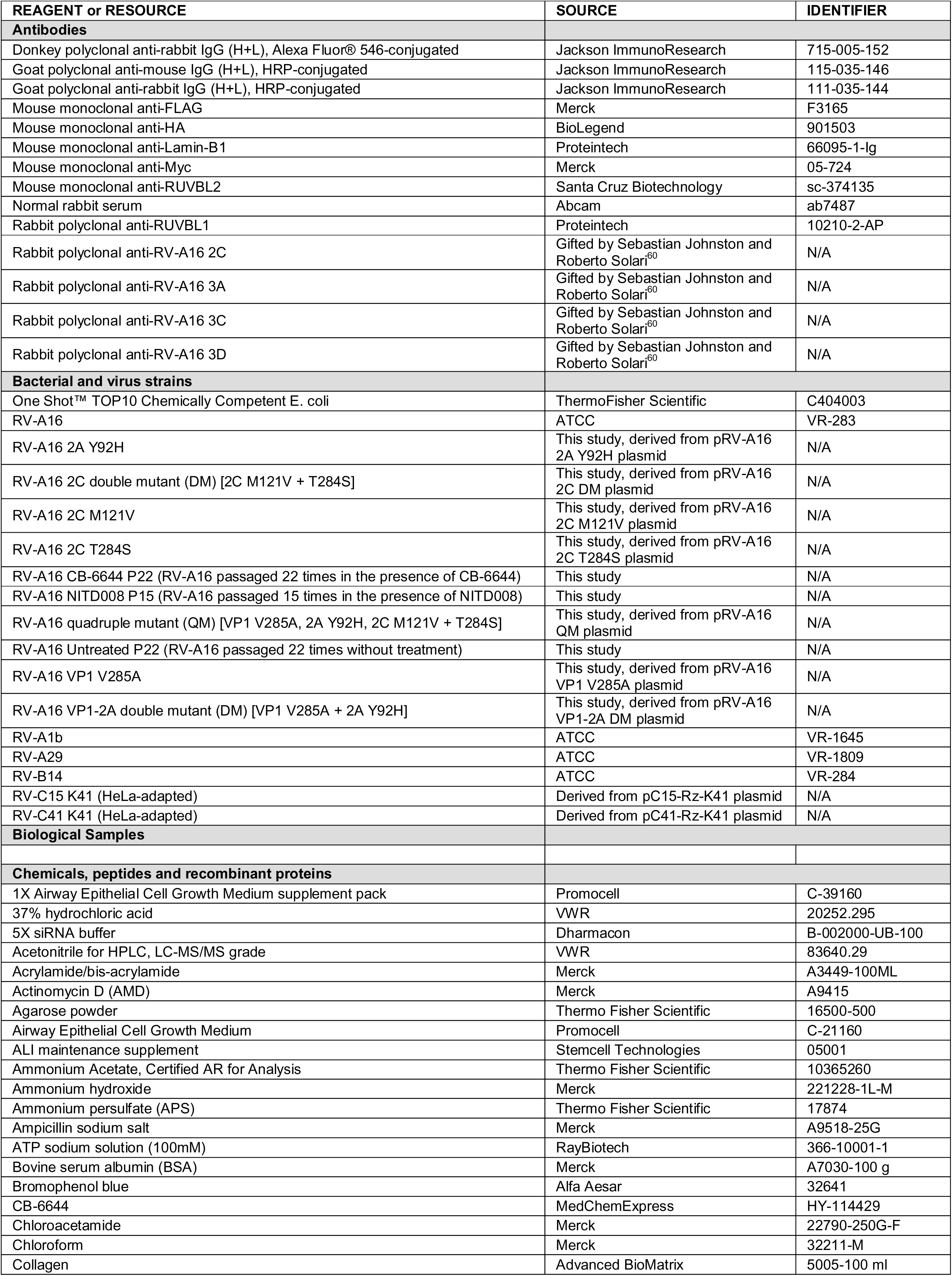
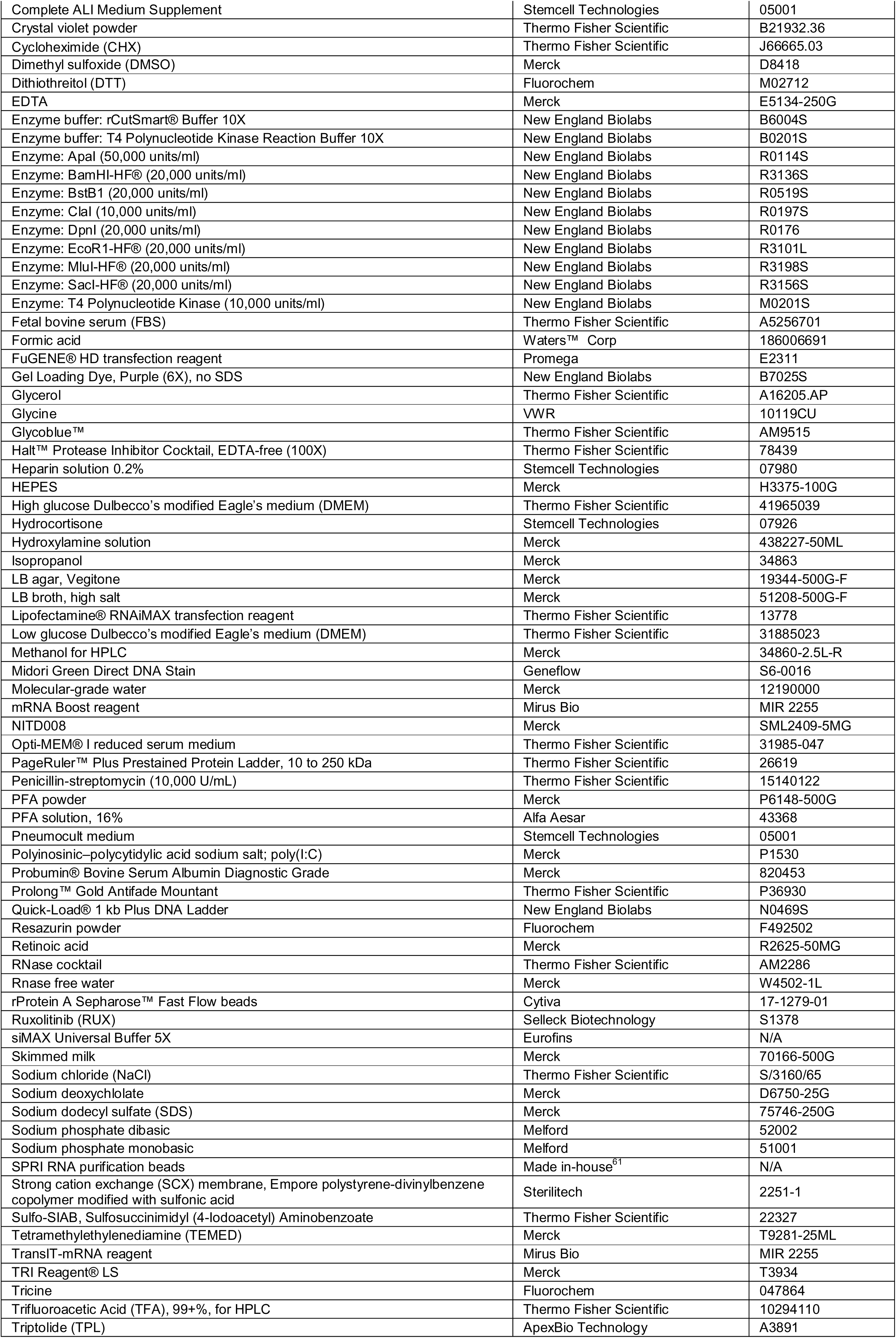
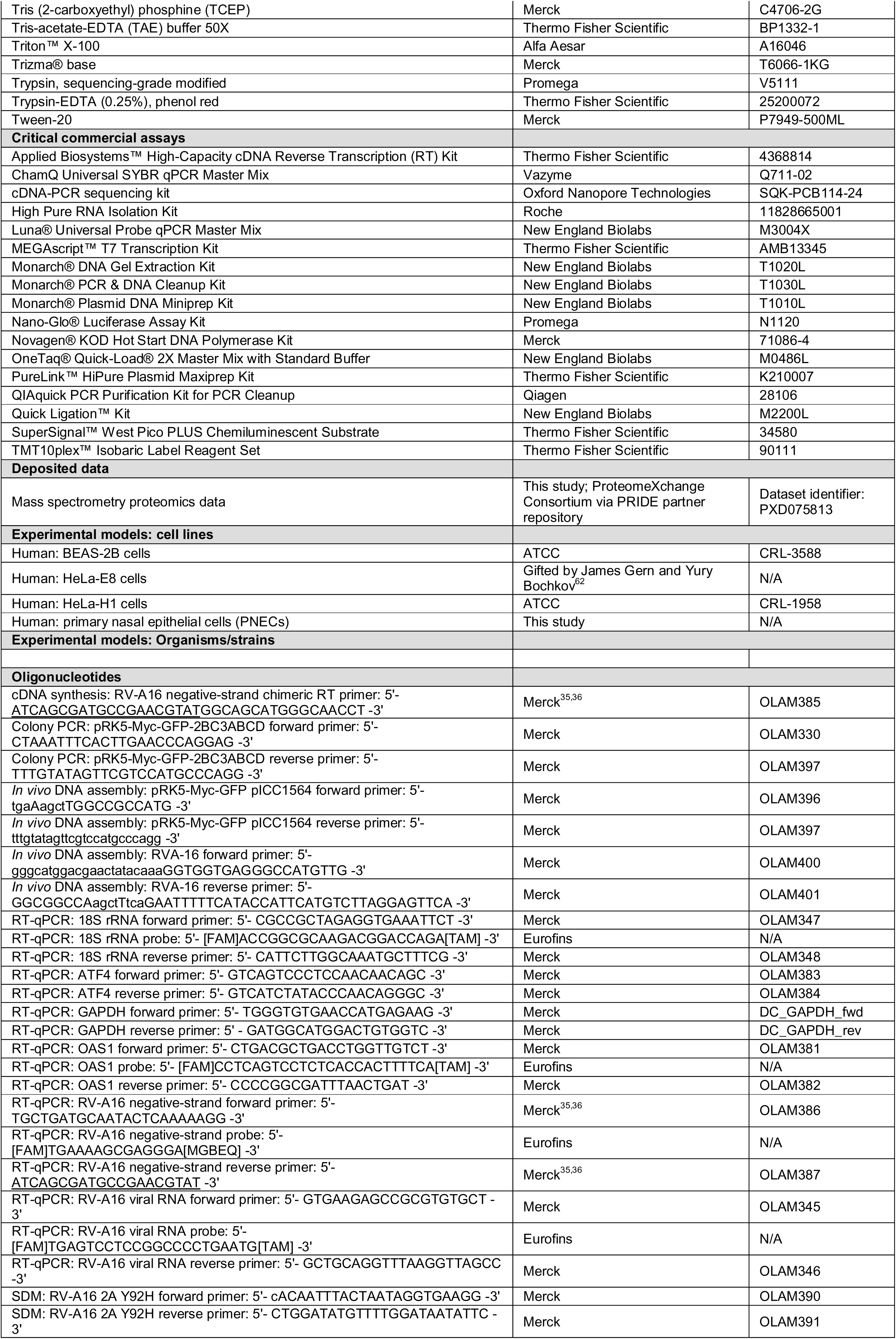
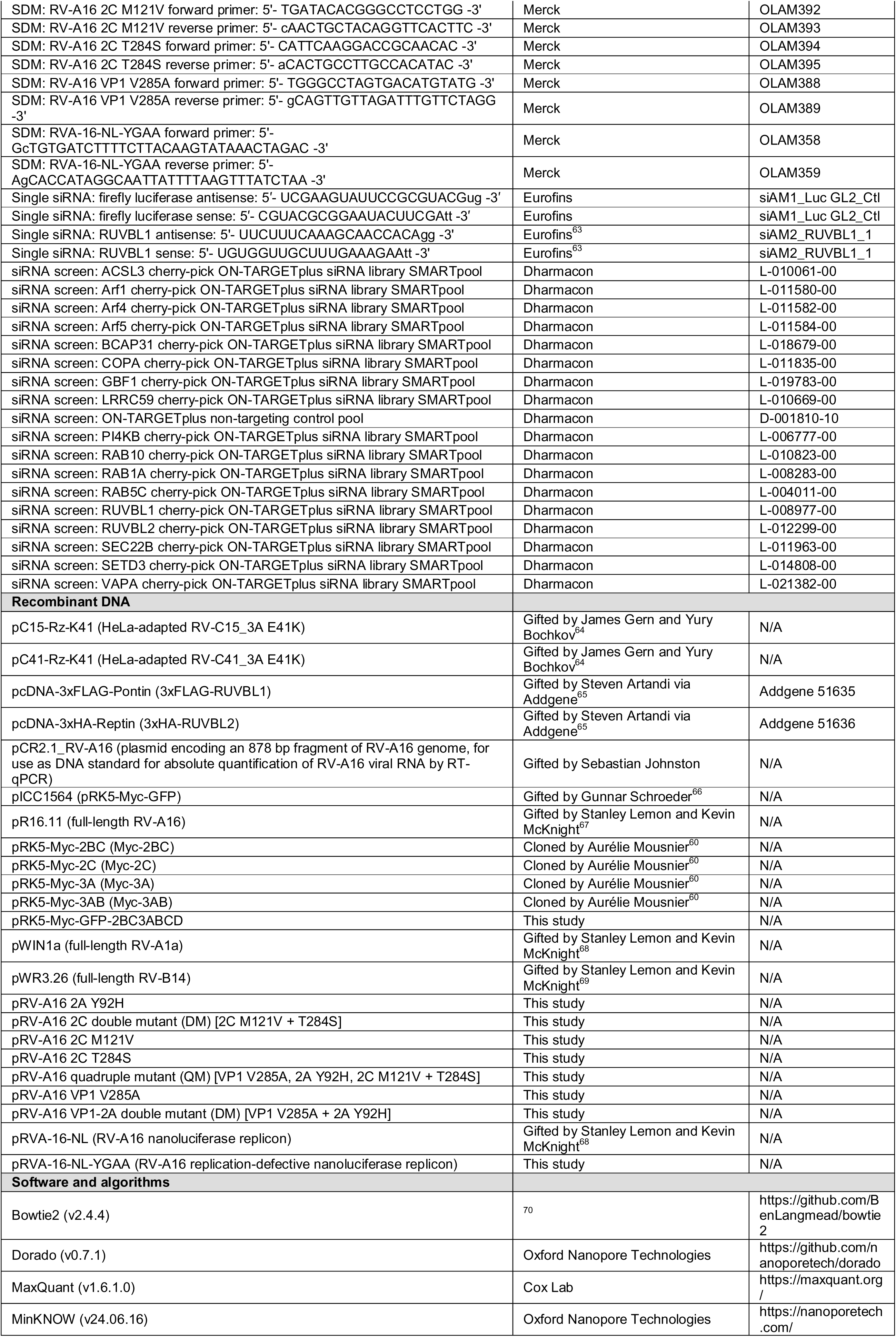
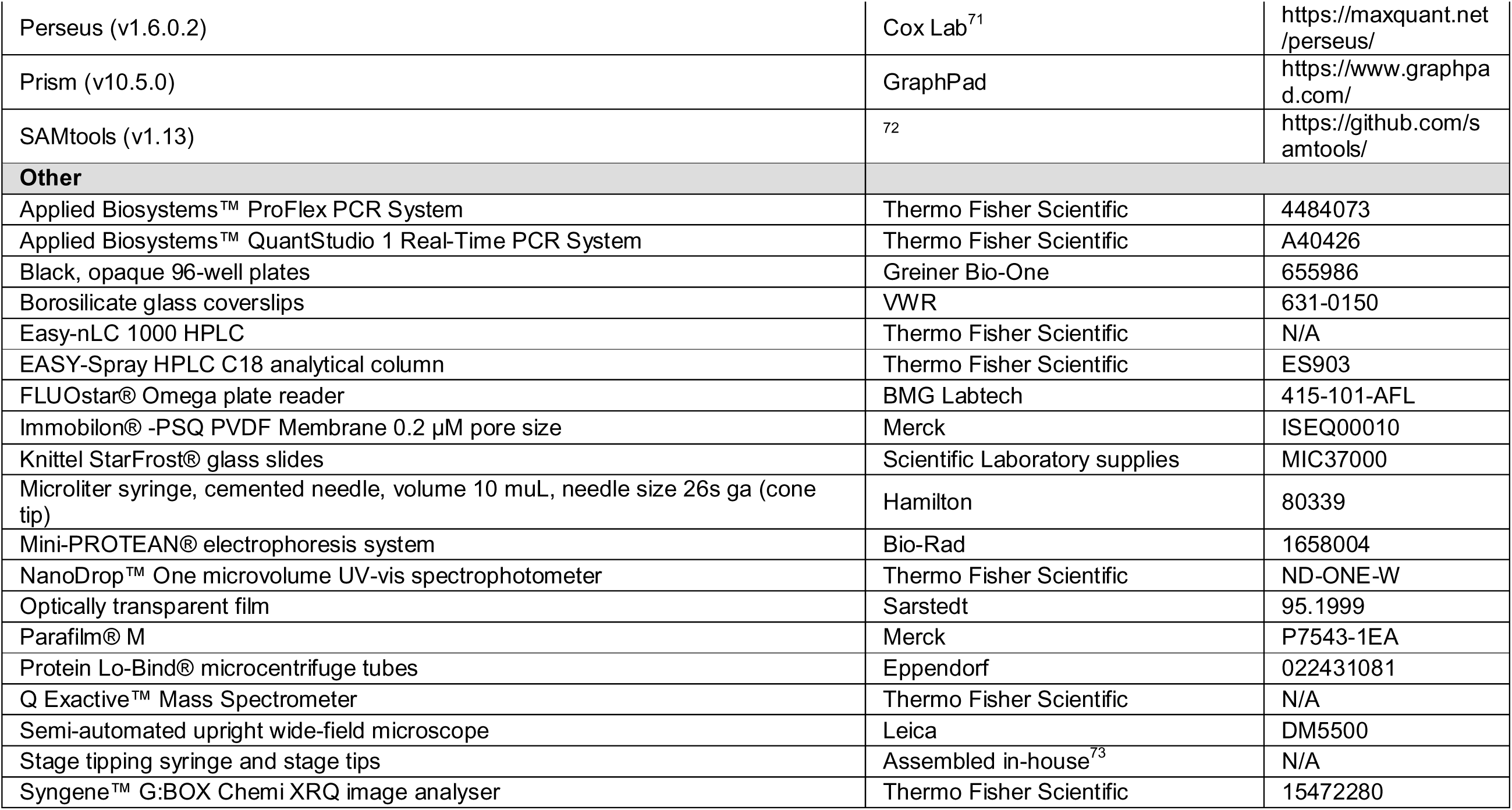

### 2 Experimental model and study participant details

#### 2.1 Cell lines

Human bronchial epithelial BEAS-2B cells (CRL-3588, ATCC) and human cervical carcinoma HeLa cells (-E8,^62^; -H1, CRL-1958, ATCC) were cultured as monolayers in high glucose Dulbecco’s modified Eagle medium (DMEM; 41965039, Thermo Fisher Scientific) supplemented with 10% (v/v) fetal bovine serum (FBS; A5256701, Thermo Fisher Scientific) (complete medium). HeLa-E8 cells were a gift from James Gern and Yury Bochkov (University of Wisconsin-Madison). Cells were maintained at 37 °C in a humidified atmosphere containing 5% CO_2_.

#### 2.2 Primary cells

##### 2.2.1 Sampling PNECs and monolayer culture

Human primary nasal epithelial cells (PNECs) were sampled, cultured and differentiated as described previously.^74,75^ All tissue culture flasks and transwells were collagen-coated (5005-100 ml, Advanced BioMatrix) overnight at 37 °C prior to use. PNECs were obtained via nasal brushing from five consenting adult volunteers (1 male, 4 female) with no known history of lung disease. After sampling (5 turns within 1 nostril), nasal brushes were placed inside 15 ml conical tubes and washed with Airway Epithelial Cell Growth Medium (C-21160, Promocell) supplemented with 1X Airway Epithelial Cell Growth Medium supplement pack (C-39160, Promocell) and 1% (v/v) penicillin/streptomycin (15140122, Thermo Fisher Scientific) (monolayer medium). Cells were centrifuged (92 x g for 5 minutes), resuspended in 2 ml monolayer medium, and transferred to a 25 cm^2^ cell culture flask placed on its side (to reduce the surface area). PNECs were maintained at 37 °C in a humidified atmosphere containing 5% CO_2_, with the monolayer medium being changed every 2-3 days. Upon reaching 80% confluence, PNECs were split to a new 25 cm^2^ flask (using the bottom surface of the flask) and then to a 75 cm^2^ flask.

##### 2.2.2 Freezing down PNECs

After expanding up to a 75 cm^2^ cell culture flask at passage 2, PNECs were frozen down for long-term storage. Briefly, PNECs were trypsinised (25200072, Thermo Fisher Scientific), resuspended in low glucose DMEM (31885023, Thermo Fisher Scientific) supplemented with 5% (v/v) FBS, and centrifuged (92 x g for 5 minutes). PNECs were resuspended in 2 ml monolayer medium and a cell count performed. The 2 ml of cells was then topped up with monolayer medium supplemented with 10% (v/v) dimethyl sulfoxide (DMSO; D8418, Merck) and 10% (v/v) FBS to have 8.0x10^5^ cells/ml. Cells were aliquoted by 1 ml into cryovials, frozen at -80 °C within a Mr Frosty™, and then transferred to -150 °C for long-term storage.

##### 2.2.3 Differentiation of PNECs – submerged phase

PNECs were thawed, centrifuged (92 x g for 5 minutes), resuspended in 1 ml monolayer medium, and transferred to a 75 cm^2^ cell culture flask containing 14 ml monolayer medium. At 100% confluence, PNECs were trypsinised, resuspended in low glucose DMEM supplemented with 5% (v/v) FBS, and centrifuged (92 x g for 5 minutes). PNECs were resuspended in 2 ml Promocell air-liquid interface (ALI) medium and a cell count performed. Promocell ALI medium was prepared as follows:

i. 500 ml Airway Epithelial Cell Growth Medium (C-21160, Promocell) was supplemented with 2X supplement packs (C-39160, Promocell) (excluding triiodothyronine and retinoic acid), 1% (v/v) penicillin/streptomycin and 3 µg/ml bovine serum albumin (BSA; A7030-100 g, Merck) (Promocell 2X medium).
ii. 500 ml low glucose DMEM (31885023, Thermo Fisher Scientific) was supplemented with 1% (v/v) penicillin/streptomycin.
iii. 25 ml Promocell 2X medium was added to 25 ml low glucose DMEM (from step ii) and supplemented with 50 µl 15 µg/ml retinoic acid (R2625-50MG, Merck) to make 50 ml Promocell ALI medium. Aliquot was protected from light.

The 2 ml of cell suspension was then topped up with Promocell ALI medium to have 3.0x10^4^ cells/250 µl. Transwells were seeded apically (250 µl/transwell), while 750 µl Promocell ALI medium was added to the basolateral compartment. To ensure adherence, cells were left undisturbed for 2 days. The apical (250 µl) and basolateral (750 µl) media were changed every 2-3 days until PNECs achieved 100% confluence (after approximately 4-8 days; cells exhibited a cobblestone appearance).

##### 2.2.4 Differentiation of PNECs – ALI phase

When PNECs reached 100% confluence, the apical and basolateral medium was completely removed, and the basolateral medium replaced with 500 µl Pneumocult ALI medium. Pneumocult ALI medium was prepared as follows:

i. 500 ml Pneumocult medium (05001, Stemcell Technologies) was supplemented with 50 ml Complete ALI Medium Supplement (05001, Stemcell Technologies) and 1% (v/v) penicillin/streptomycin.
ii. 50 ml Pneumocult medium (from step i) was supplemented with 250 µl hydrocortisone (07926, Stemcell Technologies), 100 µl 0.2% (v/v) heparin solution (07980, Stemcell Technologies) and 500 µl ALI maintenance supplement (05001, Stemcell Technologies) to make 50 ml Pneumocult ALI medium. Aliquot was protected from light.

After PNEC cultures were put into ALI, basolateral medium was changed every 2-3 days, and the cultures were closely observed for signs of mucous and beating cilia. After at least 28 days of incubation, and confirmation of the presence of mucous and extensive coverage with beating cilia, the cultures were deemed well-differentiated PNECs (WD-PNECs).

#### 2.3 Viruses

RV-A1b (VR-1645, ATCC), RV-A16 (VR-283, ATCC), RV-A29 (VR-1809, ATCC), and RV-B14 (VR-284, ATCC) were propagated in HeLa-H1 cells. Plasmids encoding full-length, HeLa-adapted RV-C15 (pC15-Rz-K41) and RV-C41 (pC41-Rz-K41) were a gift from James Gern and Yury Bochkov (University of Wisconsin-Madison).^64^ To propagate RV-C15 and RV-C41, plasmids were linearised, and viral RNA was *in vitro* transcribed and purified as described in section 3.20.12. Viral RNA was then transfected into HeLa-E8 cells to produce RV-C master stocks as described in section 3.11.

##### 2.3.1 Generation of experimental virus stocks from master stocks

Experimental virus stocks were generated as follows. Master stocks were thawed at room temperature and diluted in high glucose DMEM supplemented with 2% (v/v) FBS and 25 mM HEPES (pH 7.2-7.5) (H3375-100G, Merck) (infection medium). 2X T175 flasks of HeLa-H1 or -E8 cells were infected with 20 ml diluted virus (MOI=3) and incubated at 33 °C, 5% CO_2_ for approximately 48 hours (RV-C) or until 100% of cells exhibited cytopathic effect (CPE; RV-C does not produce observable CPE) (RV-A, -B). All RV infections in this study were performed at 33 °C to replicate the physiological conditions of the upper respiratory tract, the predominant site of RV infection *in vivo.*^76^ After three cycles of freezing and thawing to release viral particles, cell lysates were clarified by centrifugation (2,300 x g for 30 minutes at 4 °C), and supernatants were subsequently diluted 1:10 in infection medium (40 ml supernatant in 360 ml infection medium). 20X T175 flasks of HeLa-H1 or -E8 cells were then infected with 20 ml diluted virus, and incubated as above for approximately 48 hours or until 100% of cells exhibited CPE. After three cycles of freezing and thawing, cell lysates were clarified as above, and supernatants were aliquoted and transferred to -80 °C for long-term storage. 1 aliquot was kept for analysis for virus titre by 50% tissue culture infection dose (TCID_50_) assay (RV-A, -B) or plaque assay (RV-C).

### 3 Method details

#### 3.1 Determination of the virus titre of experimental samples and virus stocks

Experimental samples (cells and infection medium) were immediately frozen at -80 °C at the indicated time points. After three cycles of freezing and thawing, samples were harvested by scraping the well or dish with a sterile tip or cell scraper, and the whole cell lysate was collected. Experimental samples and virus stock aliquots (which were also freeze-thawed, see section 2.3.1) were subsequently centrifuged (15,000 x g for 15 minutes).

##### 3.1.1 TCID_50_ assay (RV-A, RV-B)

Clarified supernatants were diluted 1:10 in high glucose DMEM supplemented with 2% (v/v) FBS, 25 mM HEPES (pH 7.2-7.5) and 1% (v/v) penicillin/streptomycin (titration medium) in a 96-well plate (20 µl sample in 180 µl titration medium). For all RV-A and RV-B virus stocks and experimental samples, 4-6 technical replicates were performed for each sample. 8X 10-fold dilutions (10^-1^-10^-8^) were subsequently prepared in titration medium for each replicate, and 50 µl of each dilution was transferred to a new flat-bottomed 96-well plate. Plates were then seeded with 150 µl HeLa-H1 cells (diluted in titration medium, 1x10^5^ cells/ml) to have a final volume of 200 µl/well, and were incubated at 33 °C, 5% CO_2_ for 5 days. Plates were then manually observed for CPE under a light microscope. Wells were scored as positive if cells exhibited signs of CPE (gaps in confluence, cell rounding etc.). For each sample, the TCID_50_/ml was calculated via the Reed and Muench method.^77^

##### 3.1.2 Plaque assay (RV-C)

HeLa-E8 cells were seeded in complete medium into 12-well plates to reach 90-95% confluence on the day of plaque assay. Clarified supernatants were diluted 1:10 in infection medium in a 96-well plate (25 µl sample in 225 µl infection medium). For all RV-C virus stocks and experimental samples, 1 technical replicate was performed for each sample. 6X 10-fold dilutions (10^-1^-10^-6^) were subsequently prepared in infection medium for each replicate.

The complete medium was then removed from the HeLa-E8 cells and replaced with 200 µl of each dilution of each sample (starting from 10^-6^ and working up to 10^-1^). Plates were incubated for 1 h at 33 °C, 5% CO_2,_ and were rocked every 10 minutes to ensure even distribution of samples.

Following the 1 h incubation, the plates were removed from the incubator, and the overlay was prepared as follows. For each well to be overlayed, agarose (16500-500, Thermo Fisher Scientific) was dissolved in 750 µl infection medium to have a final concentration of 1.6% (w/v) (overlay medium). Overlay medium was then boiled in a microwave (2-3 minutes at high power) and placed in a 56 °C water bath to prevent it from solidifying. For each well to be overlayed, 750 µl warm overlay medium was then mixed with 750 µl room temperature infection medium to prepare 1.5 ml overlay (to prevent the overlay from solidifying before pouring, overlay was prepared in batches of 40 ml, with the overlay medium being placed back in the 56 °C water bath between batches). 1.5 ml overlay was then added to each well and left to solidify (at least 20 minutes at room temperature). Plates were then incubated for 5 days at 33 °C, 5% CO_2_ to allow plaques to develop.

Following the 5-day incubation, cells were fixed by adding 1 ml 4% (w/v) paraformaldehyde (PFA; P6148-500G, Merck) (dissolved in PBS) to each well and incubating for 30 minutes at room temperature. The PFA was then aspirated, and the agarose plugs were removed by placing each well under hot running water for 5 seconds. Cells were then stained by adding 1 ml 1% (w/v) crystal violet (B21932.36, Thermo Fisher Scientific) (dissolved in deionised water + 20% (v/v) methanol) to each well and incubating for 20 minutes at room temperature. Crystal violet was then aspirated, and the wells were thoroughly washed X5 in distilled water and dried overnight on tissue paper. Plaques were counted the next day as follows. For each sample, the lowest dilution with a countable number of plaques (i.e. not too many plaques or overlapping plaques) was first identified. Plaques were then counted for this dilution and all higher dilutions containing plaques. The average number of plaques across the total number of wells counted was then calculated to obtain the PFU/0.2 ml, which was subsequently multiplied by 5 to obtain the PFU/ml.

#### 3.2 RV Infection

##### 3.2.1 MOI calculation and virus dilution

All cell lines were infected with RV at 95-100% confluence at the indicated multiplicity of infection (MOI). The volume of virus stock required to achieve the desired MOI was calculated as follows. For RV-A and RV-B, the TCID_50_/ml was first converted to the number of infectious virus particles/ml (plaque forming units/ml; PFU/ml). 1 TCID_50_ = 0.7 PFU based on the Poisson distribution applied to viral infection, therefore the PFU/ml was calculated by multiplying the TCID_50_/ml by 0.7.^78^ RV-C stocks were titred by plaque assay and already expressed as PFU/ml; conversion was therefore not required.

In 1X 75 cm^2^ cell culture flask, there is approximately 13x10^6^ HeLa cells when 100% confluent, so 13x10^6^/75 = 1.73x10^5^ cells/cm^2^. This was used to extrapolate the number of cells in the wells/dishes based on their surface area and confluence. The number of cells was multiplied by the desired MOI, which was subsequently divided by the PFU/ml of the virus stock to obtain the volume (ml) of virus stock required. For each well/dish to be infected, virus stocks were diluted in infection medium to a final volume based on the manufacturer’s recommendations.

##### 3.2.2 Infection of cell lines with RV

Complete medium was removed from each well/dish and diluted virus (in infection medium) added. Infections were synchronised by incubating for 1 hour at 33 °C, 5% CO_2_ (virus adsorption period), followed by 5X washes in high glucose DMEM. Upon removal of the final wash, fresh infection medium was added. For 0-hour timepoints, infected cells were immediately processed for the required readout. For all other timepoints, infected cells were re-incubated at 33 °C, 5% CO_2_ for the required amount of time, and subsequently processed for the required readout.

##### 3.2.3 Infection of WD-PNEC cultures with RV

WD-PNECs were first meticulously examined under a light microscope. Only the highest quality cultures (i.e. those with numerous beating cilia, with similar amounts of mucous coverage, and without holes or contamination) were selected for experimentation. In well-differentiated primary airway epithelial cell (WD-PAEC) models, RVs have been shown to replicate in a small number of ciliated cells on the apical surface and to be released apically.^34,35^ For all WD-PNEC experiments in this study, cultures were therefore infected and treated apically, and virus was harvested from infected cultures via apical wash.

For each transwell to be infected, the required volume of virus stock to achieve an MOI of 0.01 (based on an average of 3.0x10^5^ cells per transwell) was diluted in 100 µl low glucose DMEM. If present, pre-treatment was removed from the apical compartment, taking care not to damage the culture. 100 µl of diluted virus was then carefully transferred to the apical compartment, with the pipette tip touching the side to minimise damage to the culture. This was denoted as time = 0 h. Infections were synchronised by incubating for 6 hours at 33 °C, 5% CO_2_ (virus adsorption period). The virus inoculum was then removed, and WD-PNECs were washed apically 6X in 300 µl low glucose DMEM. After each wash was added, WD-PNECs were incubated for 2 minutes at room temperature before removal. To confirm that no residual virus particles remained after washing, wash 6 was kept for analysis by TCID_50_ assay or plaque assay (6 h timepoint). Following the washes, the 500 µl Pneumocult ALI medium in the basolateral compartment was replaced with fresh medium. WD-PNECs were then incubated at 33 °C, 5% CO_2_.

At the required timepoints, the virus released from WD-PNEC cultures was harvested via 1X apical wash. Low glucose DMEM (280 µl if 20 µl apical treatment was present, 300 µl if no apical treatment was present) was added, and WD-PNECs were incubated for 2 minutes at room temperature. The 300 µl wash was then gently pipetted up and down, removed, and kept for analysis by TCID_50_ assay or plaque assay. Following each timepoint, the basolateral medium was changed, and the WD-PNECs were incubated at 33 °C, 5% CO_2_.

#### 3.3 General protocol for treatment of adherent cells with inhibitors for RV infection assays

The following protocol relates to data shown in Figures 2A-C, 3A-C and P, 4A and D, S1D, and S2B-D. HeLa or BEAS-2B cells were seeded in complete medium into multi-well plates. Where indicated, cells were pre-treated at 95-100% confluence with the indicated concentration of the indicated small molecule inhibitor or DMSO diluted in complete medium, for the indicated time at 37 °C, 5% CO_2_. HeLa cells were subsequently infected at 95-100% confluence with RV at the indicated MOI, alone or in the presence of the indicated small molecule inhibitor or DMSO. Immediately after virus adsorption (0 h) and/or at the indicated time point, cells were treated with the indicated small molecule inhibitor(s) or DMSO diluted in infection medium. Where indicated, uninfected untreated controls were also performed. At the indicated time points, cells were processed for the indicated readout (TCID_50_ assay, plaque assay, Western Blotting or RT-qPCR). 1 technical replicate was performed per condition per experiment.

#### 3.4 CB-6644 and NITD008 dose effect assays

HeLa cells were seeded in complete medium into multi-well plates and subsequently infected at 95-100% confluence with RV at the indicated MOI. Immediately after virus adsorption (0 h) or at the indicated time point, cells were treated with increasing concentrations of the indicated small molecule inhibitor or DMSO (diluted in infection medium) corresponding to the amount present in the highest inhibitor treatment. At 0 h and at the indicated end point, cell lysates were analysed for virus titre by TCID_50_ assay or plaque assay. 1-2 technical replicates were performed per condition per experiment. For each experiment, technical replicates for drug-treated conditions at the end point were averaged and plotted as individual points. The technical replicates for the 0 h (input) condition and DMSO-treated condition at the end point were also averaged for each experiment, and then averaged across all independent experiments and plotted as lines on each graph. IC_50_, IC_90_ and IC_99_ values were calculated as described in section 4.1.1.

#### 3.5 CB-6644 time of addition assay

HeLa-H1 cells were seeded in complete medium into multi-well plates and subsequently infected at 95-100% confluence with RV-A16 (MOI=20). Immediately after virus adsorption (0 h) or at the indicated time point, cells were treated with 500 nM CB-6644 (HY-114429, MedChemExpress) or DMSO diluted in infection medium. At 6 h, cell lysates were analysed for virus titre by TCID_50_ assay. 1 technical replicate was performed per condition per experiment.

#### 3.6 Cell viability assay for small molecule inhibitor experiments in cell lines

The following protocol relates to data shown in Figures 1G, 3L, and S2E. All cell viability assays were performed in parallel to corresponding infection or transfection experiments. Incubations were performed so that cells were exposed to the same treatments for the same amount of time as in the corresponding infection experiments (including pre-treatment time, virus adsorption time, and time after adsorption) or transfection experiments. HeLa-H1 cells were seeded in 100 µl complete medium into black, opaque 96-well plates with a transparent bottom (655986, Greiner Bio-One). At 95-100% confluence, cells were treated with the indicated small molecule inhibitor(s) at the indicated concentration(s) (the highest concentration(s) used in the infection or transfection experiment) or DMSO diluted in complete or infection medium. Cells were subsequently incubated with the infection or transfection plate at 37 °C or 33 °C (depending on the experiment), 5% CO_2_ for the indicated amount of time.

At the endpoint, complete or infection medium only was added to X5-6 wells as blanks. 1 ml 1.5 mg/ml resazurin stock (F492502, Fluorochem) was diluted in 9 ml PBS to make a working solution of 0.15 mg/ml, which was vortexed well and protected from light. 20 µl 0.15 mg/ml resazurin was then added to the 100 µl complete or infection medium within each well, and plates were incubated for 1 hour at 37 °C, 5% CO_2_. Fluorescence was then recorded using a 560 nm excitation / 590 nm emission filter set on a FLUOstar® Omega plate reader (415-101-AFL, BMG Labtech). Raw fluorescence intensity values were normalised based on the average of the X5-6 blank wells. For each experiment, 3-6 technical replicates were performed for drug-treated conditions, 5-6 were performed for the DMSO-treated condition, and 5-6 were performed for blank wells.

#### 3.7 WD-PNEC experiments

##### 3.7.1 Prophylactic assay

X2 WD-PNEC cultures were used per treatment (CB-6644 or DMSO) per donor. Data are representative of 3 (RV-A16 and RV-B14) or 2 (RV-C15) independent biological donors. WD-PNEC cultures were pre-treated apically with 20 µl CB-6644 or DMSO diluted in low glucose DMEM. Cultures were incubated overnight for 16 h at 37°C, 5% CO_2_.

Following overnight incubation, pre-treatments were removed, and cultures were infected apically with RV as described in section 3.2.3 in the presence of the corresponding treatment. Following apical washes at 6 h, cultures were treated apically with 20 µl CB-6644 or DMSO diluted in low glucose DMEM, and were incubated at 33°C, 5% CO_2_. Following the single apical wash (280 µl) at all subsequent timepoints (24, 48, 72, 96, 120, 144 and 168 h), cultures were re-treated with 20 µl CB-6644 or DMSO.

After three cycles of freezing and thawing, apical washes were analysed for virus titre by TCID_50_ assay (RV-A, -B) or plaque assay (RV-C). For each treatment at each timepoint, the TCID_50_/ml (RV-A, -B) or PFU/ml (RV-C) of X2 WD-PNEC cultures was averaged and plotted as a single point on each graph.

##### 3.7.2 Therapeutic assay

X2 WD-PNEC cultures were used per treatment (time of CB-6644 or DMSO addition) per donor. Data are representative of 2 independent biological donors. WD-PNEC cultures were infected apically with RV-A16 as described in section 3.2.3 in the absence of any treatment. Following apical washes at 6 h, X2 cultures were treated with 1.5 µM CB-6644 and X2 cultures were treated with DMSO, both diluted in 20 µl low glucose DMEM (6 h treatment). The remainder of the cultures received no treatment, and all cultures were subsequently incubated at 33°C, 5% CO_2_.

Following the single apical wash at 12 h (280 µl for cultures that were treated at 6 h, 300 µl for all remaining cultures), cultures that were treated at 6 h were re-treated with 1.5 µM CB-6644 or DMSO. In addition, X2 cultures not previously treated were treated with 1.5 µM CB-6644 (12 h treatment). The remainder of the cultures received no treatment, and all cultures were subsequently re-incubated at 33°C, 5% CO_2_. This process was repeated after the single apical wash at the 24 h and 36 h timepoints, with cultures that were previously treated being re-treated with CB-6644 or DMSO, and X2 cultures receiving their first CB-6644 treatment. At all subsequent timepoints (48, 72, 96, 120, 144 and 168 h), all cultures were re-treated with CB-6644 or DMSO following the single apical wash.

After three cycles of freezing and thawing, apical washes were analysed for virus titre by TCID_50_ assay. For each treatment at each timepoint, the TCID_50_/ml of X2 WD-PNEC cultures was averaged and plotted as a single point on each graph.

##### 3.7.3 Cell viability assay for WD-PNEC cultures

X4 WD-PNEC cultures were used per treatment (CB-6644 or DMSO) per donor. Data are representative of 3 independent biological donors. WD-PNEC cultures were treated apically with 20 µl 2.0 µM CB-6644 or DMSO diluted in low glucose DMEM, and incubated for 192 h at 33°C, 5% CO_2_. Every 24 h, this treatment was removed and replaced with 20 µl fresh 2.0 µM CB-6644 or DMSO. At 190 h, 1X fresh WD-PNEC culture was treated apically with 100 µl 0.1% (v/v) Triton™ X-100 (A16046, Alfa Aesar) diluted in low glucose DMEM, and incubated for the remaining 2 h of the experiment at 33°C, 5% CO_2_. This well served as a positive control. At 192 h, the apical treatments were removed and 200 µl low glucose DMEM was added to each well. 200 µl low glucose DMEM also added to X2 empty transwells, which served as blank wells.

The working solution of 0.15 mg/ml resazurin was prepared as described in section 3.6. 40 µl 0.15 mg/ml resazurin were then added to 200 µl low glucose DMEM within the apical compartment of each WD-PNEC culture or empty transwell, and cultures/transwells were incubated for 2 h at 37°C, 5% CO_2_. The 240 µl low glucose DMEM within each culture/transwell were then gently pipetted up and down, and 120 µl was transferred to a flat-bottomed 96-well plate. Fluorescence was recorded on a plate reader as described in section 3.6. Raw fluorescence intensity values were normalised based on the average of the X2 blank wells.

#### 3.8 Transfection of cell lines

##### 3.8.1 Reverse transfection of siRNA pools (siRNA screen)

siRNA pools for screening each of the 17 LC-MS/MS hits were purchased from Dharmacon (Cherry-pick ON-TARGETplus siRNA library SMARTpool). Each pool targeted a single gene and was comprised of a set of 4 siRNA. Details of these siRNA pools are shown in the key resources table. ON-TARGETplus non-targeting control pool was also purchased from Dharmacon (D-001810-10). Each siRNA pool was resuspended in 1X siRNA buffer (B-002000-UB-100, Dharmacon), aliquoted, and stored at -80 °C according to manufacturer’s instructions. siRNA pools were reverse-transfected into HeLa-H1 cells as follows. For each well to be transfected, 4 pmol siRNA pool was diluted in 20 µl Opti-MEM® I reduced serum medium (31985-047, Thermo Fisher Scientific). 0.3 µl Lipofectamine® RNAiMAX transfection reagent (13778, Thermo Fisher Scientific) (vortexed and warmed to room temperature) was then added, and the transfection mix was mixed gently, transferred to 1X well of a 96-well plate, and incubated for 30 minutes at room temperature. 80 µl HeLa-H1 cells (diluted in complete medium) was subsequently seeded on top of the transfection mix to have a final siRNA concentration of 40 nM. Transfected cells were then incubated for 72 h at 37 °C, 5% CO_2_ until 95-100% confluent.

Following the 72 h incubation, transfected cells were infected with RV-A16 (MOI=20). At 6 h, cell lysates were analysed for virus titre by TCID_50_ assay. For each of the 17 siRNA pools targeting LC-MS/MS hits, X1 technical replicate was performed per experiment. For the non-targeting control pool, X6 technical replicates were performed per experiment. For each experiment, the mean TCID_50_/ml of the X6 non-targeting control pool wells was first calculated. The mean TCID_50_/ml of the non-targeting control pool across the 3-4 independent experiments (mean of means) was then calculated. For each experiment, the TCID_50_/ml for each of the 17 siRNA pools targeting LC-MS/MS hits was calculated as percentage of the mean of means of the non-targeting control pool.

##### 3.8.2 Cell viability assay for siRNA screen

Of the 17 siRNA pools that were investigated in the siRNA screen, only the 9 that produced a statistically significant reduction in RV-A16 titres were carried forward for cell viability assay. siRNA pools were reverse-transfected into HeLa-H1 cells as described above, except reactions were prepared in black, opaque 96-well plates with a transparent bottom (655986, Greiner Bio-One).

Following the 72 h incubation, 100 µl complete medium only was added to X6 wells as blanks. The working solution of 0.15 mg/ml resazurin was prepared as described in section 3.6. 20 µl 0.15 mg/ml resazurin was then added to the 100 µl complete medium within each well, and plates were incubated for 1 hour at 37 °C, 5% CO_2_. Fluorescence was recorded on a plate reader as described in section 3.6. Raw fluorescence intensity values were normalised based on the average of the X6 blank wells. For each experiment, X3 technical replicates were performed for each of the 9 siRNA pools targeting the LC-MS/MS hits, X6 were performed for the non-targeting control pool, and X6 were performed for blank wells. For each experiment, the mean fluorescence intensity of the X6 non-targeting control pool wells was first calculated. The mean fluorescence intensity of the non-targeting control pool across the 3 independent experiments (mean of means) was then calculated. For each experiment, the mean fluorescence intensity for each of the 9 siRNA pools targeting LC-MS/MS hits (X3 wells each) was then calculated as a percentage of the mean of means of the non-targeting control pool.

##### 3.8.3 Forward transfection of single siRNA

The following protocol relates to data shown in Figure 3E-J, S1B-C, and S1E-F. Single siRNA targeting RUVBL1^63^ and firefly luciferase were purchased from Eurofins. Details of these siRNA are shown in the key resources table. Each siRNA was resuspended in 1X siMAX Universal Buffer (included with siRNA, Eurofins), aliquoted, and stored at -80 °C according to manufacturer’s instructions. siRNA were forward-transfected into HeLa-H1 cells as follows. HeLa-H1 cells were seeded in complete medium into 24-well plates (directly onto the well surface or onto glass coverslips) so to reach 30% confluence on the day of transfection. For each well to be transfected, 5 pmol siRNA was diluted in 50 µl Opti-MEM® I and, separately, 0.5 µl Lipofectamine® RNAiMAX (vortexed and warmed to room temperature) was diluted in 50 µl OptiMEM® I. Both mixtures were incubated for 5 minutes at room temperature, before being combined, mixed gently, and incubated for a further 20 minutes at room temperature. The complete medium was then removed from each well and replaced with 400 µl high glucose DMEM supplemented with 1% (v/v) FBS (low serum medium). Each well was then topped up with 100 µl transfection mix to have a final siRNA concentration of 10 nM. Plates were gently rocked back and forth to mix, and subsequently incubated for 48 h at 37 °C, 5% CO_2_. Low serum medium was then removed from each well and replaced with 500 µl complete medium, and plates were incubated for a further 24 h at 37 °C, 5% CO_2_.

Following the 72 h incubation, transfected cells were either immediately processed for the required readout, or were infected with RV-A16 (MOI=20) and then processed for the required readout(s) at the indicated timepoint(s). 1 technical replicate was performed per condition per experiment.

##### 3.8.4 Forward transfection of plasmid DNA

Details of all plasmids used are shown in the key resources table. Plasmid DNA was transfected into HeLa-H1 cells as follows. HeLa-H1 cells were seeded in complete medium into multi-well plates or dishes to reach 90-95% confluence on the day of transfection. For each well or dish to be transfected, transfection mixes were prepared at a 3:1 ratio of FuGENE® HD transfection reagent (E2311, Promega) (µl) to DNA quantity (µg) (see below table). If multiple plasmids were transfected into the same well or dish, the total quantity of DNA was split between each plasmid as required, and the corresponding amount of each plasmid was transfected.

For each well or dish to be transfected, the required quantity of plasmid DNA was first diluted in the required volume of Opti-MEM® I. The required volume of FuGENE® HD (vortexed and warmed to room temperature) was then added, and the transfection mix was immediately vortexed for 10 seconds and subsequently incubated for exactly 15 minutes at room temperature. The transfection mix was then added dropwise directly into the well or dish containing HeLa-H1 cells in complete medium. Plates or dishes were then gently rocked back and forth to mix, and were subsequently incubated at 37 °C, 5% CO_2_. At the indicated time points, medium was removed and replaced with fresh complete medium, and cells were processed at the indicated endpoint for the indicated readout. 1 technical replicate was performed per condition per experiment.

| Well/dish | DNA (µg) | FuGENE® HD (µl) | Opti-MEM® I to a total volume of (µl) | HeLa-H1 in complete medium (µl) |
| --- | --- | --- | --- | --- |
| 24-well plate | 0.6 | 1.8 | 30 | 0.5 |
| 6-well plate | 2.92 | 8.77 | 146 | 2.0 |
| 100 mm dish | 19.12 | 57.37 | 956 | 10.0 |

##### 3.8.5 Reverse transfection of in vitro transcribed RNA

*In-vitro* transcribed RNA was transfected into HeLa cells as follows. For each well to be transfected, the required quantity of RNA was first diluted in the required volume of Opti-MEM® I (see below table). The required volumes of *Trans*IT-mRNA reagent and mRNA Boost reagent (both MIR 2255, Mirus Bio) (both vortexed and warmed to room temperature) were then added, and the transfection mix was mixed gently and incubated for exactly 5 minutes at room temperature. The required volume of HeLa cells (diluted in complete medium to reach 95-100% confluence the next day) was added on top of the transfection mix, and the total volume (transfection mix + cells) was then seeded per well of a multi-well plate. Plates were then gently rocked back and forth to mix, and were subsequently incubated at 37 °C, 5% CO_2_. At the indicated time points, cells were processed for the indicated readout. 1 technical replicate was performed per condition per experiment.

| Well | Opti-MEM®<br>I<br>(μl) | RNA<br>(μg) | TransIT-mRNA<br>reagent<br>(μl) | mRNA<br>Boost<br>reagent<br>(μl) | HeLa-H1 in<br>complete<br>medium<br>(μl) | Total<br>volume to<br>seed<br>(μl) |
| --- | --- | --- | --- | --- | --- | --- |
| 96-well plate | 10 | 0.1 | 0.2 | 0.2 | 90 | 100 |
| 24-well plate | 50 | 0.5 | 0.5 | 0.5 | 450 | 500 |
| 6-well plate | 250 | 2.5 | 5 | 5 | 2,250 | 2,500 |

#### 3.9 Bypass entry assay

Plasmids encoding full-length RV-A16 (pR16.11), RV-A1a (pWIN1a) and RV-B14 (pWR3.26) were gifts from Stanley Lemon and Kevin McKnight (University of North Carolina).^67–69^ Plasmids were linearised, and viral RNA was *in vitro* transcribed from linearised plasmids and purified as described in section 3.20.12.

Transfection mixes for *in vitro* transcribed RV-A16, RV-A1a, or RV-B14 RNA were prepared and seeded for the required number of 24-well plate wells, as described in section 3.8.5. Immediately after seeding, cells (in 500 µl) were treated with 2 µl 125 µM CB-6644 (diluted in complete medium) to have a final concentration of 500 nM, or were treated with the corresponding volume of DMSO. Plates were then gently rocked back and forth to mix, and were subsequently incubated for 14 h at 37 °C, 5% CO_2_. At 14 h, cell lysates were analysed for virus titre by TCID_50_ assay. 1 technical replicate was performed per condition per experiment.

#### 3.10 Nanoluciferase replicon assay

The plasmid encoding the replication-defective RV-A16 nanoluciferase replicon (pRVA-16-NL-YGAA) was generated by site-directed mutagenesis as described in section 3.20.9. The plasmid was then linearised, and mutant RNA was *in vitro* transcribed from the linearised plasmid and purified as described in section 3.20.12.

Transfection mixes for *in vitro* transcribed RV-A16 replication-defective nanoluciferase replicon RNA were prepared and seeded for the required number of 96-well plate wells, as described in section 3.8.5. Cells were seeded into separate 96-well plates for each timepoint (0, 6 and 24 h). Immediately after seeding, cells (in 100 µl) were treated with 2 µl 9 mM cycloheximide (CHX; J66665.03, Thermo Fisher Scientific) (freshly prepared and diluted in complete medium) to achieve a final concentration of 178 µM; with 2 µl 25 µM CB-6644 (diluted in complete medium) to achieve a final concentration of 500 nM; or with the corresponding volume of DMSO. Plates were rocked gently back and forth to mix, and the 0 h plate was immediately processed for Nano-Glo® Luciferase Assay (N1120, Promega). The 6 h and 24 h plates were incubated at 37 °C, 5% CO_2_ and processed for Nano-Glo® Luciferase Assay at the indicated endpoint.

Nano-Glo Luciferase Assay was performed as follows. Assay components and 96-well plates were first incubated at room temperature for 15 minutes. For each well to be assayed, 2 µl luciferase substrate was diluted in 98 µl assay buffer to make 100 µl assay reagent. 100 µl assay reagent was then added to each well and mixed gently, and plates were incubated for 3 minutes at room temperature while protected from light. The complete medium containing assay reagent was then gently pipetted up and down, and 100 µl from each well was transferred to a fresh white, opaque 96-well plate. Luminescence was then recorded using the top lens optic (LENS) on a FLUOstar® Omega plate reader (BMG Labtech) in luminescence mode with an integration time of 1.0 s per well and auto-gain enabled. For each experiment, X1 technical replicate was performed per timepoint per condition. For each condition (CHX, CB-6644 or DMSO) within each experiment, luminescence values at 0 h were subtracted from luminescence values at 6 h and 24 h. All 0 h-subtracted values were then normalised to the 0 h-subtracted DMSO value at 24 h.

#### 3.11 Reverse transfection of RNA for virus stock production

Transfection mixes for *in vitro* transcribed RNA were prepared and seeded for the required number of 6-well plate wells, as described in section 3.8.5. Plates were then gently rocked back and forth to mix, and were subsequently incubated at 37 °C, 5% CO_2_ for approximately 144 hours (RV-C) or until 100% of cells exhibited CPE (RV-A). After three cycles of freezing and thawing, cell lysates were clarified by centrifugation (2,300 x g for 30 minutes at 4 °C), and supernatants were subsequently diluted 1:8 in infection medium (2.5 ml supernatant in 17.5 ml infection medium). 1X T175 flasks of HeLa cells was then infected with 20 ml diluted virus and incubated at 33°C, 5% CO_2_ for approximately 48 hours (RV-C) or until 100% of cells exhibited CPE (RV-A). After three cycles of freezing and thawing, the supernatants were clarified as described in section 2.3.1, aliquoted, and transferred to -80 °C for long-term storage. 1 aliquot was kept for analysis for virus titre by TCID_50_ assay or plaque assay.

#### 3.12 Poly(I:C) stimulation assay

HeLa-H1 cells were seeded in complete medium into 48-well plates. At 95-100% confluence, 0 h wells were immediately processed for RNA extraction. For 10 h wells, complete medium was removed and replaced with 2 µg/ml actinomycin D (AMD; A9415, Merck), 10 µM triptolide (TPL; A3891, ApexBio Technology), or 10 µM ruxolitinib (RUX; S1378, Selleck Biotechnology) (all diluted in infection medium), or with infection medium alone, and incubated for 1 h at 33°C, 5% CO_2_. These treatments were then removed and replaced with 50 µg/ml polyinosinic–polycytidylic acid sodium salt (poly(I:C); P1530, Merck) alone or in combination with AMD, TPL, or RUX (all diluted in infection medium), or with infection medium alone, and plates were incubated for a further 9 h at 33 °C, 5% CO_2_. At 10 h, wells were processed for RNA extraction. OAS1 mRNA levels at 0 h and 10 h were quantified by RT-qPCR, normalised to 18S ribosomal RNA (rRNA) levels, and then expressed relative to OAS1 mRNA levels at 0 h. 1 technical replicate was performed per condition per experiment.

#### 3.13 Polyprotein processing assay (transfection)

The plasmid encoding Myc-GFP-2BC3ABCD (pRK5-Myc-GFP-2BC3ABCD) was generated by *in vivo* DNA assembly as described in section 3.20.10.

HeLa-H1 cells were seeded in complete medium into 6-well plates. At 95-100% confluence, cells were transfected with pRK5-Myc-GFP-2BC3ABCD as described in section 3.8.4, or were left untransfected. Immediately after transfection, cells (in 2,146 µl) were treated with 10.7 µl 100 µM CB-6644 (diluted in complete medium) to achieve a final concentration of 500 nM; with the corresponding volume of DMSO; or were left untreated. Plates were rocked gently back and forth to mix, centrifuged (210 x g for 5 minutes), and incubated for 4 h at 37 °C, 5% CO_2_. Treatments were subsequently removed and replaced with fresh 500 nM CB-6644 or DMSO (diluted in complete medium), or with complete medium alone, and plates were incubated for a further 17 h at 37 °C, 5% CO_2_. At 21 h, cell lysates were analysed for Myc and RV-A16 2C/2BC, 3C/3CD and 3A/3AB by Western Blotting.

In parallel, HeLa-H1 cells were seeded in complete medium into 6-well plates and subsequently infected at 95-100% confluence with RV-A16 (MOI=20), or were left uninfected. At 8 h, cell lysates were analysed for RV-A16 2C/2BC, 3C/3CD and 3A/3AB by Western Blotting. 1 technical replicate was performed per condition per experiment.

#### 3.14 SDS-PAGE and Western Blotting

##### 3.14.1 Sample preparation and quantification

For data shown in Figures 3B-C, F-H and O, S1B-E, and S3A, at the indicated timepoints, complete or infection medium was removed, and cells were washed 1X in PBS. PBS was then completely removed, and cells were lysed as follows. 200 µl fresh 1 M dithiothreitol (DTT; M02712, Flurochem) was first added to 800 µl 2X Laemmli sample buffer comprised of 0.15 M Tris-HCl (pH=6.8) (Trizma® base, T6066-1KG, Merck; hydrochloric acid, 20252.295, VWR), 25% (v/v) glycerol (A16205.AP, Thermo Fisher Scientific), 5% (w/v) sodium dodecyl sulfate (SDS; 75746-250G, Merck) and RNase free water (W4502-1L, Merck). 2X Laemmli sample buffer with 0.2 M DTT was then vortexed and warmed for 5 minutes at 100 °C. The required volume was then added to each well/dish, cells were harvested using a cell scraper, and the whole cell lysate was collected. Samples were then vortexed for 10 seconds, centrifuged (15,000 x g for 5 minutes at room temperature), and subsequently incubated for 5 minutes at 100 °C. This 3-step process was repeated until samples were non-viscous. For samples for Figures 3B-C and F-H, and S1B-E, samples were then quantified using the dsDNA setting on a NanoDrop™ One microvolume UV-vis spectrophotometer (ND-ONE-W, Thermo Fisher Scientific). The volume (µl) of each sample containing the required quantity of DNA (µg) to load per well was calculated, and this volume was transferred to a fresh microcentrifuge tube. Samples were then topped up to equivalent volumes with 2X Laemmli sample buffer with 0.2 M DTT. Samples for Figures 3O and S3A were not quantified, as the HeLa-H1 cells in each well were of highly similar confluence at the time of cell lysis. 1 µl bromophenol blue (32641, Alfa Aesar) was added to all samples. Immediately prior to loading, samples were vortexed for 10 seconds, incubated for 5 minutes at 100 °C, and centrifuged (15,000 x g for 5 minutes at room temperature). Equivalent volumes were loaded per well.

For immunoprecipitation (IP; Figure S3B-E) and co-immunoprecipitation (co-IP; Figures 4E and S4A-J) experiments, cells were lysed in the indicated buffer as described in sections 3.18.1 and 3.18.3. Samples were then processed for IP (sections 3.18.1 and 3.18.3) or co-IP (sections 3.18.1 and 3.18.2), and the input and pull-down fractions were collected for analysis by Western Blotting. All input fractions were diluted with 5X Laemmli sample buffer (0.3 M Tris-HCl pH=6.8, 50% (v/v) glycerol, 10% (w/v) SDS, RNase free water and 0.5 M DTT) to have a final Laemmli sample buffer concentration of 1X. For co-IPs, pull-down fractions were eluted in 2X Laemmli sample buffer with DTT, as described in section 3.18.2. For IP of NSPs for polyprotein processing assay, pull-down fractions were eluted in 2X Laemmli sample buffer without DTT, and DTT was added after elution as described in section 3.18.3. Samples for IPs and co-IPs were not quantified. 1 µl bromophenol blue was added to all input and pull-down fractions. Immediately prior to loading, samples were vortexed for 10 seconds, incubated for 5 minutes at 100 °C, and centrifuged (15,000 x g for 5 minutes at room temperature). Equivalent volumes were loaded per well.

##### 3.14.2 SDS-PAGE

SDS-polyacrylamide resolving gels were prepared according to Bio-Rad manuals with the following composition: either 8% (v/v) or 12% (v/v) acrylamide/bis-acrylamide (A3449-100ML, Merck), 375 mM Tris-HCl (pH=8.8), 0.1% SDS (w/v), 0.05% (w/v) ammonium persulfate (APS; 17874, Thermo Fisher Scientific), 0.1% (v/v) tetramethylethylenediamine (TEMED; T9281-25ML, Merck), and RNase free water. Resolving gels with 15% (v/v) acrylamide were prepared with the same composition, but with a lower TEMED concentration of 0.05% (v/v). SDS-polyacrylamide stacking gels were prepared with the following composition: 4% (v/v) acrylamide/bis-acrylamide, 125 mM Tris-HCl (pH=6.8), 0.1% (w/v) SDS, 0.05% (w/v) APS, 0.2% (v/v) TEMED, and RNase free water. SDS-PAGE was performed using the Mini-PROTEAN® electrophoresis system (1658004, Bio-Rad). Gels were assembled in the gel tank on ice. For all gels except those for RV-A16 3C, 3A and 3AB, gels were run in standard SDS-PAGE running buffer composed of 0.19 M glycine (10119CU, VWR), 25 mM Tris, and 0.1% (w/v) SDS. For gels for RV-A16 3C, 3A and 3AB, gels were run using separate anode and cathode buffers. Anode buffer (100 mM Tris, pH=8.9) was added to the outer compartment, and cathode buffer (100 mM Tris, 100 mM Tricine (047864, Fluorochem), 0.1% (w/v) SDS, pH=8.3) was added to the inner compartment. Gels were electrophoresed at 60 V until samples left the stacking gel, and then at 100 V until protein resolution was achieved. Samples were run in parallel to PageRuler™ Plus Prestained Protein Ladder, 10 to 250 kDa (26619, Thermo Fisher Scientific).

##### 3.14.3 Western Blotting

SDS-PAGE gels were assembled with sponges, filter paper, and Immobilon® -PSQ PVDF Membrane (ISEQ00010, Merck) to form a transfer sandwich, which was subsequently placed inside a wet electrophoretic transfer system (Mini Trans-Blot, Bio-Rad) on ice. Tanks were filled with transfer buffer (0.19 M glycine, 25 mM Tris), and transfer was performed at 100 V for 20-60 minutes according to protein size. If membranes were to be probed for multiple targets, membranes were then cut horizontally as required. Membranes were subsequently washed in TBS buffer (20 mM Tris, 150 mM sodium chloride (NaCl; S/3160/65, Thermo Fisher Scientific), pH=7.4-7.6) supplemented with 0.1% (v/v) Tween-20 (P7949-500ML, Merck) (TBS-Tween) (1X 30 second wash) and then blocked in 3% (w/v) skimmed milk (70166-500G, Merck) in TBS-Tween for 1 hour at room temperature on a tube roller. Membranes were briefly washed in TBS-Tween (1X 30 second wash) prior to the addition of primary antibodies/antisera. Primary antibodies/antisera were diluted in 3% (w/v) skimmed milk (in TBS-Tween) and incubated with membranes overnight at 4 °C on a tube roller. Following the overnight incubation, membranes were washed thoroughly in TBS-Tween at room temperature (2X 30 second washes, 3X 5 minute washes) prior to the addition of secondary antibodies. Secondary antibodies were diluted in 3% (w/v) skimmed milk (in TBS-Tween) and incubated with membranes for 1 hour at room temperature on a tube roller. Following the 1 hour incubation, membranes were washed thoroughly in TBS-Tween at room temperature (2X 30 second washes, 3X 5 minute washes). 1 ml SuperSignal™ West Pico PLUS Chemiluminescent Substrate (34580, Thermo Fisher Scientific) was then added to each membrane, and protein bands were revealed using a Syngene™ G:BOX Chemi XRQ image analyser (15472280, Thermo Fisher Scientific). See key resources table for full list of all antibodies and antisera used for Western Blotting. Antisera against RV-A16 2C, 3A, 3C and 3D were gifts from Sebastian Johnston and Roberto Solari (Imperial College London).^60^

##### 3.14.4 Densitometry

For Figures 3B-C and F-H, and S1B-E, densitometric analysis of Western Blot membranes was performed using ImageJ software (NIH). Horizontal rectangles were drawn around all the bands of interest for a given protein, and the bands were subsequently quantified to generate peaks. The background was then removed from each peak to determine the specific intensity value for each band. For RV-A16 3C, all 3 visible bands were quantified and compounded for each lane. For each lane, the intensity value of the band(s) for the protein of interest was normalised to the intensity value of the band for the loading control (e.g. Lamin-B1). The normalised intensity value of the band(s) for the protein of interest in the treated condition (e.g. drug-treated) was then calculated as a percentage of the normalised intensity value of the band(s) for the protein of interest in the control condition (e.g. DMSO-treated).

#### 3.15 Immunofluorescence (RUVBL1 knockdown assay)

HeLa-H1 cells were seeded in complete medium into 24-well plates containing borosilicate glass coverslips (631-0150, VWR) to reach 30% confluence on the day of transfection. Cells were then transfected with a single siRNA targeting RUVBL1 or firefly luciferase as described in section 3.8.3, and incubated for 72 h at 37 °C, 5% CO_2_. Following the 72 h incubation, transfected cells were infected with RV-A16 (MOI=20), or were left uninfected. At 6 h, adherent cells were fixed and stained for immunofluorescence as follows (all washes and incubations inside 24-well plates were performed at room temperature and in 500 µl). Infection medium was removed from each well, and cells were washed in high glucose DMEM (3X 30 seconds) and subsequently fixed by incubating in 3% (v/v) PFA (43368, Alfa Aesar) (diluted in PBS, 1X 15 minutes). Cells were then washed in PBS (3X 30 seconds), and were either immediately stained or temporarily stored in PBS at 4 °C.

PBS was then removed from each well, and residual PFA was quenched by incubating cells in 0.1 M glycine (dissolved in PBS, 2X 10 minutes). Cells were then washed in PBS (3X 30 seconds), and were subsequently permeabilised by incubating in 0.1% (v/v) Triton™ X-100 (in PBS, 1X 10 minutes). Cells were then washed in PBS (3X 30 second washes) and were subsequently blocked by incubating in 5% (v/v) FBS (in PBS, 1X 30 minutes).

Primary antiserum (targeting RV-A16 2C) (see key resources table for full list of all antibodies and antisera used for immunofluorescence) were diluted in PBS and distributed onto Parafilm® M (P7543-1EA, Merck) in 30 µl drops. Coverslips were then transferred on top of the drops (cell-side down) and incubated in a humid atmosphere for 1 hour at room temperature. Coverslips were then transferred back into 24-well plate wells (cell-side up) and washed thoroughly in PBS (1X 30 seconds, then 2X 5 minutes). Secondary antibodies were diluted in PBS and distributed onto Parafilm® M in 30 µl drops. Coverslips were then transferred on top of the drops (cell-side down) and incubated in a humid atmosphere and protected from light for 1 hour at room temperature. Coverslips were then transferred back into 24-well plate wells (cell-side up) and washed thoroughly in PBS (1X 30 seconds, then 2X 5 minutes). Coverslips were then washed briefly in distilled water, drained completely, and mounted onto glass slides (Knittel StarFrost® via Scientific Laboratory supplies, MIC37000) in 10 µl Prolong™ Gold Antifade Mountant (P36930, Thermo Fisher Scientific). Mounted coverslips were protected from light and left to cure overnight at room temperature, and were subsequently stored briefly at 4 °C. All imaging was performed within 24 hours of staining on a semi-automated upright wide-field microscope (DM5500, Leica). 1 technical replicate was performed per condition per experiment, and 5 images were captured per condition. For each image, the number of RV-A16 2C-positive cells was manually counted and expressed as a percentage of the total number of cells. The percentage of RV-A16 2C-positive cells was then averaged across all 5 images per condition.

#### 3.16 RT-qPCR

##### 3.16.1 RNA extraction from cell lines

At the indicated timepoints, HeLa-H1 cells in multi-well plates were washed in PBS (1X 30 second wash at room temperature). Total RNA was subsequently extracted from each well using the High Pure RNA Isolation Kit (11828665001, Roche) according to manufacturer’s instructions, except for the cell lysis step. For each well, 400 µl Lysis/Binding Buffer (11828665001, Roche) was diluted in 200 µl D-PBS, and 600 µl diluted Lysis/Binding Buffer was then added per well. Samples were harvested by scraping the well with an RNase-free tip, and the whole cell lysate was collected and vortexed for 15 seconds. Each sample was then transferred to a High Pure Filter Tube (11828665001, Roche), and the protocol was continued according to manufacturer’s instructions. The eluted, purified RNA for each sample was quantified using the RNA setting on a NanoDrop™ One microvolume UV-vis spectrophotometer (ND-ONE-W, Thermo Fisher Scientific), and subsequently stored at -80 °C.

##### 3.16.2 cDNA synthesis

Purified RNA was reverse-transcribed for complementary DNA (cDNA) synthesis using the Applied Biosystems™ High-Capacity cDNA Reverse Transcription (RT) Kit (4368814, Thermo Fisher Scientific). For samples for data shown in Figure 3E, 250 ng RNA was reverse transcribed to cDNA (these experiments were performed in 24-well plates). For all other samples, 200 ng RNA was reverse transcribed to cDNA (these experiments were performed in 48-well plates, and as a result a smaller total quantity of RNA was extracted per sample).

For each sample, the required volume of purified RNA (containing 250 ng or 200 ng) was transferred to the appropriate number of PCR tubes (X1 tube per cDNA reaction) and topped up to 10 µl with RNase-free water (W4502-1L, Merck). For each sample, a general 2X RT master mix was prepared according to manufacturer’s instructions (10X RT buffer, 25X dNTP mix, 10X random primers, RT enzyme, and RNase-free water) for synthesis of random-primed cDNA (for analysis by RT-qPCR for RV-A16 viral RNA, 18S rRNA, OAS1, ATF4 or GAPDH). Note that ‘viral RNA’ refers to total RV-A16 RNA, the vast majority of which during infection is comprised of positive-strand RNA.^35,36^

To specifically quantify RV-A16 negative-strand RNA, an alternative approach to cDNA synthesis was required. As with other positive-strand RNA viruses, RV-A16 positive-strand RNA can “self-prime” during RT, even in the absence of primers. This self-primed positive-strand RNA can be subsequently amplified by conventional RT-qPCR targeting negative-strand RNA, resulting in a loss of negative-strand specificity.^35,36,79–81^ To circumvent this, we utilised a chimeric RT primer which extends RV-A16 negative-strand RNA at the 5’ end with a heterologous sequence derived from the genome of *Escherichia coli*.^35,36^ This sequence was later used to selectively amplify negative-strand RNA by RT-qPCR (see section 3.16.3). For each sample for the negative-strand RNA timecourse (Figure 3P), one 2X RT master mix was prepared as described above for synthesis of random-primed cDNA (for analysis of 18S rRNA), and a second 2X RT master mix was prepared in which the random primers were substituted with 10 µM chimeric primer (final concentration 1 µM) for synthesis of negative-strand-specific cDNA (for analysis of negative-strand RNA). Details of the chimeric RT primer are shown in the key resources table.

For all 2X RT master mixes, a 2X no RT control master mix was also prepared, in which RT enzyme was replaced with RNAse-free water. 10 µl 2X master mix was then added to each diluted RNA sample to have a total volume of 20 µl containing 1X master mix. RT was then performed on an Applied Biosystems™ ProFlex PCR System (4484073, Thermo Fisher Scientific) using the protocol described below. To remove contaminating RNA, all newly synthesised negative-strand-specific cDNA samples were subsequently treated with 1 µl RNase cocktail (AM2286, Thermo Fisher Scientific), incubated for 1 h at 37 °C, purified using the Monarch® PCR & DNA Cleanup Kit (T1030L, New England Biolabs) according to manufacturer’s instructions, and eluted in 20 µl RNase free water. All synthesised cDNA samples were subsequently stored at -20 °C.

| Step | Temperature (°C) | Time (minutes) | Number of cycles |
| --- | --- | --- | --- |
| 1 | 25 | 10:00 | X1 |
| 2 | 37 | 120:00 | X1 |
| 3 | 85 | 5:00 | X1 |
| 4 | 4 | Hold | X1 |

##### 3.16.3 RT-qPCR

For analysis of all genes except ATF4 and GAPDH (Figure S2D), RT-qPCR reaction mixes were prepared using Luna® Universal Probe qPCR Master Mix (M3004X, New England Biolabs). For analysis of ATF4 and GAPDH, RT-qPCR reaction mixes were prepared using ChamQ Universal SYBR qPCR Master Mix (Q711-02, Vazyme). All RT-qPCR reaction mixes were prepared according to manufacturer’s instructions.

For analysis of RV-A16 viral RNA, RT-qPCR reaction mixes were prepared using a final concentration of 0.2 µM probe, 0.05 µM forward primer and 0.3 µM reverse primer. For analysis of all other genes except ATF4 and GAPDH, reaction mixes were prepared using a final concentration of 0.2 µM probe, 0.4 µM forward primer and 0.4 µM reverse primer. Note that to ensure specificity for negative-strand RNA, the reverse primer for analysis of negative-strand RNA anneals to the heterologous *E.coli* sequence incorporated into the cDNA, as described in section 3.16.2.^35,36^ For analysis of ATF4 and GAPDH, reaction mixes were prepared without a probe and using a final concentration of 0.4 µM forward primer and 0.4 µM reverse primer. Details of all primers and probes used for RT-qPCR are shown in the key resources table.

RT-qPCR reaction mixes were aliquoted into white, opaque PCR plates (72.1979.010, Sarstedt), and 12.5 ng of the appropriate cDNA was added to the appropriate wells (for cDNA made from 200 ng RNA, 1.25 µl 10 ng/µl cDNA was added per well; for cDNA made from 250 ng RNA, 1 µl 10 ng/µl cDNA was added per well). For each RT-qPCR plate, a no RT control for each gene being analysed was assembled alongside the experimental samples. For all RT-qPCR plates analysing RV-A16 viral RNA, a set of 10 DNA standards of known concentration (pCR2.1_RV-A16 serially diluted in RNase free water from 1x10^10^ to 1x10^1^ copies/µl) was also assembled alongside the experimental samples to generate a standard curve. pCR2.1_RV-A16 was a gift from Sebastian Johnston (Imperial College London). Each experimental sample was run in duplicate; each of the 10 DNA standards and each no RT control was run in singlet. Plates were sealed with optically transparent film (95.1999, Sarstedt), and reactions were cycled on an Applied Biosystems™ QuantStudio 1 Real-Time PCR System (A40426, Thermo Fisher Scientific) as described below.

| Step | Temperature | Time (seconds) | Number of cycles |
| --- | --- | --- | --- |
| 1 | 95 | 00:60 | X1 |
| 2 | 95 | 00:15 | X40 |
| 3 | 60 | 00:30 (+ plate read) |  |

For RV-A16 viral RNA, the exact number of copies in each of the 2 replicates for each sample was extrapolated from a standard curve generated by amplification of the 10 DNA standards. The average number of copies in each sample was then calculated and normalised to the level of 18S rRNA (according to the average Cq for 18S in each sample). For cDNA made from 200 ng RNA, the average number of copies was multiplied by 5 to determine the average number of copies/µg of RNA. For cDNA made from 250 ng RNA, the average number of copies was multiplied by 4 to determine the average number of copies/µg of RNA.

For RV-A16 negative-sense RNA and OAS1, the average Cq of the 2 replicates for each sample was normalised to the level of 18S rRNA (according to the average Cq for 18S in each sample), and expressed as fold-change relative to the 0 h timepoint. For ATF4, the average Cq of the 2 replicates for each sample was normalised to the level of GAPDH (according to the average Cq for GAPDH in each sample) and expressed as fold-change relative to the untreated uninfected condition.

#### 3.17 Negative-sense RNA timecourse

HeLa-H1 cells were seeded in complete medium into 48-well plates (1 plate per timepoint) and subsequently infected at 95-100% confluence with RV-A16 (MOI=20). Immediately after virus adsorption (0 h), cells were treated with infection medium, and the 0 h plate was processed for RNA extraction. All other plates were incubated for 1 h at 33 °C, 5% CO_2_. Infection medium was subsequently removed from all wells and replaced with 500 nM CB-6644 or DMSO (diluted in infection medium), and incubated at 33 °C, 5% CO_2_. At the indicated timepoints, cells were processed for RNA extraction. RV-A16 negative-sense RNA levels at 0 h and at the indicated timepoints were quantified by RT-qPCR, normalised to 18S ribosomal RNA (rRNA) levels, and then expressed relative to RV-A16 negative-sense RNA levels at 0 h. 1 technical replicate was performed per condition per experiment.

#### 3.18 Immunoprecipitation

##### 3.18.1 General immunoprecipitation protocol

HeLa-H1 cells were seeded into 100 mm or 150 mm dishes. At 95-100% confluence, cells were processed for infection or transfection as required. At the indicated time points, infection or complete medium was removed and dishes were washed 1X in PBS. To preserve weak and/or transient protein-protein interactions, cells were then cross-linked as follows. All PBS was removed, and cells were treated with 1% (v/v) PFA (in PBS) (43368, Alfa Aesar) (5 ml per 100 mm dish, 8 ml per 150 mm dish) for 20 minutes at room temperature. Without removing the PFA, 1.25 M glycine (10119CU, VWR) was then added to each dish to quench the reaction (500 µl per 100 mm dish, 800 µl per 150 mm dish). Dishes were rocked back and forth to mix, incubated for 5 minutes at room temperature, and subsequently washed 3X in PBS. After the final wash, absolutely all remaining PBS was removed, and dishes were transferred to a constant temperature room (4 °C) for cell lysis.

To lyse cross-linked HeLa-H1 cells, cells were treated with 1 ml (per 100 mm dish and 150 mm dish, except where indicated) ice-cold radioimmunoprecipitation (RIPA) buffer (pH = 7.6), composed of 0.1 M sodium phosphate (monobasic, 51001, Melford; dibasic, 52002, Melford), 150 mM NaCl, 1% (v/v) Triton™ X-100, 2 mM EDTA (E5134-250G, Merck), 0.5% (w/v) sodium deoxycholate (D6750-25G, Merck) and 0.5% (w/v) SDS. RIPA buffer was supplemented with 1X Halt™ Protease Inhibitor Cocktail (78439, Thermo Fisher Scientific). Dishes were then incubated for 30 minutes at 4 °C on a rocking platform. Dishes were then scraped, and cell lysates were collected, vortexed for 10 seconds, and centrifuged (15,000 x g for 10 minutes at 4 °C). Clarified supernatants were then transferred to fresh microcentrifuge tubes, and an aliquot corresponding to 10% of the total volume of each sample was collected for analysis by Western Blotting (IP input). The remainder of the clarified supernatant was processed for IP as follows. The appropriate volume of the required antiserum or antibody was added to each sample, and samples were incubated for 2 h at 4 °C on a rotating wheel. Clarified supernatants were subsequently transferred to fresh microcentrifuge tubes containing 10 µl rProtein A Sepharose™ Fast Flow beads (17-1279-01, Cytiva), and incubated overnight at 4 °C on a rotating wheel.

Following the overnight incubation, samples for co-IP were processed as described in section 3.18.2. Samples for mass spectrometry were processed as described in section 3.19.1.

##### 3.18.2 Co-immunoprecipitation assay

HeLa-H1 cells were seeded in complete medium into 100 mm dishes to reach 90-95% confluence on the day of transfection. Cells were subsequently transfected with plasmids encoding Myc-tagged RV NSPs (pRK5-Myc-2C, -2BC, -3A or -3AB)^60^ or Myc-GFP (pICC1564) as described in section 3.8.4. pICC1564 was a gift from Gunnar Schroeder (Queen’s University Belfast).^66^ These plasmids were co-transfected with either 3xFLAG-RUVBL1 (pcDNA-3xFLAG-Pontin) or 3xHA-RUVBL2 (pcDNA-3xHA-Reptin) (double transfection; Figure S4C-D and F-J) or with both 3xFLAG-RUVBL1 and 3xHA-RUVBL2 (triple transfection; Figures 4E and S4A-B and E). pcDNA-3xFLAG-Pontin (51635, Addgene) and pcDNA-3xHA-Reptin (51636, Addgene) were gifted as bacterial stab cultures by Steven Artandi (Stanford University).^65^ Dishes were gently rocked back and forth to mix, and were subsequently incubated at 37 °C, 5% CO_2_. At 30 h post-transfection, cells were cross-linked and processed for IP using rProtein A Sepharose™ Fast Flow beads as described in section 3.18.1. IP was performed using anti-Myc (05-724, Merck) (Figures 4E and S4A-I) or anti-FLAG (F3165, Merck) (Figure S4J) antibody.

Following the overnight incubation with antibody and clarified supernatant, beads were pelleted by centrifugation (79 x g for 2 minutes at 4 °C), and the flow-through was removed. Beads were subsequently washed 20X in RIPA buffer without protease inhibitor cocktail (for each wash, beads were resuspended in 1 ml RIPA buffer without protease inhibitor cocktail, incubated for 2 minutes at 4 °C on a rotating wheel, and then pelleted by centrifugation at 79 x g for 2 minutes at 4 °C). After the final wash, the residual RIPA buffer was completely removed from the beads using a microlitre syringe (80339, Hamilton). 40 µl elution buffer (2X Laemmli sample buffer supplemented with DTT) was added to the dry beads, and beads were vortexed for 10 seconds and incubated for 20 minutes at 100 °C. Beads were then centrifuged (15,000 x g for 1 minute at room temperature), and the pull-down fraction was collected using a microlitre syringe. Input and pull-down fractions were analysed by Western Blotting for Myc and FLAG and/or HA (901503, BioLegend). 1 technical replicate was performed per condition per experiment.

##### 3.18.3 Immunoprecipitation of NSPs for polyprotein processing assay (infection)

The protocol for IP of NSPs for polyprotein processing assay (Figure S3B-E) differed significantly from the protocol for IP of NSPs for mass spectrometry (Figure 1A, section 3.18.1) and the protocol for co-IP assay (Figures 4E and S4A-J, sections 3.18.1 and 3.18.2). HeLa-H1 cells were seeded in complete medium into 150 mm dishes. X64 dishes were seeded in total (X28 for the infected, 500 nM CB-6644 condition; X28 for the infected, 5 µM NITD008 condition; X4 for the infected only condition; and X4 for the uninfected condition). To minimise elution of IgG heavy and light chains, anti-RV-A16 NSP antisera was crosslinked to rProtein A Sepharose™ Fast Flow beads 24 h prior to infection as follows. 10 µl rProtein A Sepharose™ Fast Flow beads were used per condition. For all subsequent washes, beads were resuspended in the indicated volume of buffer, incubated for 2 minutes at room temperature on a rotating wheel, and then pelleted by centrifugation. All centrifugation steps were performed at 79 x g for 2 minutes at room temperature. Beads were first washed 1X in 1 ml 0.05% (v/v) Tween-20 in PBS (PBS-Tween), and were then blocked by incubation in 1 ml 1% (w/v) Probumin® BSA (820453, Merck) in PBS for 1 h at room temperature on a rotating wheel. Beads were then washed 3X in PBS-Tween, and were subsequently incubated in 1 ml 2 mM Sulfo-SIAB (22327, Thermo Fisher Scientific) in PBS for 20 minutes at room temperature on a rotating wheel and protected from light. Beads were then washed 3X in PBS-Tween. After the final wash, beads were resuspended in PBS-Tween and distributed in equal volumes to fresh microcentrifuge tubes for each IP antisera. The required volume of anti-RV-A16 NSP antisera (2C, 3A, 3C or 3D) was then added to each tube, and beads were incubated overnight at room temperature on a rotating wheel and protected from light.

The following day, at 95-100% confluence, cells were infected with RV-A16 (MOI=20), or were left uninfected. Immediately after virus adsorption, cells were treated with 500 nM CB-6644 or 5 µM NITD008 (SML2409-5MG, Merck) (both diluted in infection medium), or were left untreated. At 8 h, cells were either PFA cross-linked as described in 3.18.1 and lysed in 1 ml ice-cold RIPA buffer (with the same composition as in 3.18.1, except the 0.5% (w/v) SDS was replaced with 0.1% (w/v) SDS) (Figure S3D), or were not PFA cross-linked and lysed in 1 ml ice-cold 1% (v/v) Triton™ X-100 buffer (with the same composition as the RIPA buffer described in section 3.18.1, except the SDS and sodium deoxycholate was removed) (Figure S3B-C and E). Both buffers were supplemented with 1X protease inhibitor cocktail. Dishes were incubated for 30 minutes at 4 °C on a rocking platform. Dishes were then scraped thoroughly, and the lysates for each condition were subsequently pooled together into 50 ml or 15 ml centrifuge tubes. Lysates were vortexed for 10 seconds and clarified by centrifugation (2,300 x g for 10 minutes at 4 °C). The clarified supernatants were then transferred to fresh 50 ml or 15 ml centrifuge tubes. For the experiment in which cells were not cross-linked (Figure S3B-C and E), clarified supernatants were then sonicated for 30 seconds on ice. Clarified supernatants were then pre-cleared by incubation with 10 µl rProtein A Sepharose™ Fast Flow beads for 1 hour at 4 °C on a rotating wheel. The beads were subsequently pelleted by centrifugation (2,300 x g for 2 minutes at 4 °C), and the clarified supernatants were transferred to fresh 50 ml or 15 ml centrifuge tubes. 200 µl of each clarified supernatant was kept for analysis by Western Blotting (IP input).

Following the overnight incubation, the beads were removed from the rotating wheel. For all subsequent washes, beads were resuspended in the indicated volume of buffer, incubated for 2 minutes at room temperature on a rotating wheel, and then pelleted by centrifugation. All centrifugation steps were performed at 79 x g for 2 minutes at room temperature. Beads were first washed 3X in 1 ml PBS-Tween, and non-crosslinked antibody was subsequently removed by incubation with 1 ml stripping buffer (100 mM glycine, 500 mM NaCl, pH=2.3-2.5) for 1 hour at room temperature on a rotating wheel. Beads were then washed 3X in 1 ml PBS-Tween. After the final wash, beads were resuspended in PBS-Tween and 10 µl beads was distributed to fresh 15 ml centrifuge tubes or microcentrifuge tubes (1 tube for each IP antisera for each condition). Equivalent volumes of pre-cleared, clarified supernatants were then transferred on top of the crosslinked beads, and were incubated overnight at 4 °C on a rotating wheel.

Following the overnight incubation, the beads were removed from the rotating wheel. For all subsequent washes, beads were resuspended in the indicated volume of buffer, incubated for 2 minutes at 4 °C on a rotating wheel, and then pelleted by centrifugation. All centrifugation steps were performed at 79 x g for 2 minutes at 4 °C. Beads were first pelleted by centrifugation. Beads in 15 ml centrifuge tubes were first resuspended in PBS-Tween, and transferred to microcentrifuge tubes. Beads were then washed X25 in 1 ml PBS-Tween. After the final wash, the residual PBS-Tween was completely removed from the beads using a microlitre syringe. 35 µl elution buffer (2X Laemmli sample buffer without DTT) was added to the dry beads, and beads were subsequently vortexed for 10 seconds and incubated for 20 minutes at room temperature with gentle shaking. Beads were then centrifuged (15,000 x g for 1 minute at room temperature), and the pull-down fraction was collected using a microlitre syringe for analysis by Western Blotting. 8.75 µl 1 M DTT was added to the 35 µl elution (final concentration 0.2 M). 1 technical replicate was performed per condition per experiment.

#### 3.19 Proteomics

##### 3.19.1 Sample preparation

All conditions for proteomics sample preparation (uninfected, 2C IP; infected 4.5 h, 2C IP; infected 4.5 h, control IP; infected 6 h, 2C IP; and infected 6 h, control IP) were run in triplicate (X3 dishes per condition). Protein Lo-Bind® microcentrifuge tubes (022431081, Eppendorf) were used at the appropriate stages throughout this protocol. HeLa-H1 cells were seeded into 150 mm dishes and infected at 95-100% confluence with RV-A16 (MOI=20), or were left uninfected. At 4.5 h and 6 h, cells were PFA cross-linked and processed for IP using rProtein A Sepharose™ Fast Flow beads as described above in section 3.18.1, with minor modifications. Briefly, dishes were lysed in 2 ml ice-cold RIPA buffer (supplemented with 1X protease inhibitor cocktail), and incubated for 30 minutes at 4 °C on a rocking platform. Dishes were then scraped thoroughly, and the lysates were subsequently transferred to microcentrifuge tubes, vortexed for 10 seconds, and clarified by centrifugation (15,000 x g for 10 minutes at 4 °C). The clarified supernatants for each triplicate (around 7.8 ml in total) were then pooled together and mixed gently, and equivalent volumes were re-distributed to fresh microcentrifuge tubes (6X tubes in total, 2X tubes per replicate, 1.3 ml per tube). The clarified supernatants were then incubated with the appropriate volume of anti-RV-A16 2C antiserum, or control normal rabbit serum (ab7487, Abcam), for 2 h at 4 °C on a rotating wheel. Clarified supernatants were subsequently transferred to fresh microcentrifuge tubes containing 5 µl rProtein A Sepharose™ Fast Flow beads, and incubated overnight at 4 °C on a rotating wheel.

For all subsequent washes, beads were resuspended in the indicated volume of buffer, incubated for 2 minutes at 4 °C on a rotating wheel, and then pelleted by centrifugation. All centrifugation steps were performed at 79 x g for 2 minutes at 4 °C. Following the overnight incubation, beads were pelleted by centrifugation and resuspended in 500 µl of flow-through. Replicates were then combined together into a single microcentrifuge tube to have 10 µl rProtein A Sepharose™ Fast Flow beads in total. Beads were then washed 5X in 1 ml RIPA buffer, 5X in 400 µl 50 mM HEPES (pH = 8.0, freshly prepared), and 1X in 200 µl 50 mM HEPES with 5 mM tris (2-carboxyethyl) phosphine (TCEP) (C4706-2G, Merck) and 10 mM chloroacetamide (22790-250G-F, Merck). After the final wash, beads were resuspended in 25 µl 50 mM HEPES with 5 mM TCEP and 10 mM chloroacetamide. 1 µl trypsin (V5111, Promega) was then added, and the beads were mixed gently and incubated overnight at 37 °C while shaking at 900 RPM.

##### 3.19.2 TMT labelling and multiplexing

Following the overnight incubation, the beads were centrifuged at 79 x g for 2 minutes at room temperature, and the 25 µl buffer containing tryptic peptides was carefully removed and transferred to a fresh microcentrifuge tube. Samples were then labelled using the TMT10plex™ Isobaric Label Reagent Set (90111, Thermo Fisher Scientific). The required TMT labels (in 0.08 mg aliquots) were each resuspended in 20 µl 100% (v/v) acetonitrile (83640.29, VWR) according to manufacturer’s instructions. 20 µl of each sample was then mixed with 20 µl of a specific TMT label (126, 127N, 127C, 128N, 128C, 129N, 129C, 130N, 130C or 131), and subsequently incubated for 2 h at room temperature while shaking at 900 RPM. Subsequently, 1 µl 5% (v/v) hydroxylamine (438227-50ML, Merck) (in 50 mM HEPES) was added to each sample to quench the reaction. Differently labelled samples were then combined together into pools according to technical replicate (X3 pools in total, X1 pool per replicate). Pooled samples were then incubated at 45 °C in a vacuum concentrator until all liquid was completely evaporated, and were subsequently stored at -80 °C.

##### 3.19.3 Stage tipping (6-layer fractionation)

Stage tips and stage tipping syringe were assembled according to established protocols.^73^ All stage tips were assembled using strong cation exchange (SCX) membranes (Empore polystyrene-divinylbenzene copolymer modified with sulfonic acid via Sterlitech, 2251-1). For each stage tip to be prepared, a stage tipping syringe was used to place a small (1 mm wide) piece of SCX membrane at the bottom of a 200 µl pipette tip. Care was taken to place the SCX membranes at the same distance from the bottom of each tip. Stage tips were then placed inside microcentrifuge tubes with a small hole cut in the lid. Note that for all centrifugation steps in the stage tipping protocol, care was taken to ensure that the entire buffer/sample had passed through the SCX membrane before proceeding to the next step. Except where indicated, all centrifugation steps were performed at 1,150 x g for 15 minutes at room temperature, with extra time being added if the buffer/sample had not totally passed through the membrane after the initial spin. SCX membranes were first activated by addition of 150 µl acetonitrile into each stage tip. Stage tips were then centrifuged, washed in 150 µl molecular-grade water (12190000, Merck), and centrifuged again.

The TMT-labelled and pooled samples were thawed on ice, resuspended in 150 µl 1% (v/v) trifluoroacetic acid (TFA; 10294110, Thermo Fisher Scientific) (pH = 1.0), and centrifuged at 13,000 x g for 5 minutes at room temperature. 135 µl (corresponding to 90%) of the supernatant of each sample was then transferred to a prepared stage tip (the remaining 15 µl was not used in order to avoid clogging the SCX membrane). Stage tips were then centrifuged. Samples (now bound to the SCX membranes) were subsequently desalted by washing stage tips 3X in 60 µl 0.2% (v/v) TFA, with a centrifugation step between each wash. After the final wash, each stage tip was transferred to a fresh microcentrifuge tube with a small hole cut in the lid. Each sample was then eluted in 6 fractions using 6 buffers of increasing stringency. Buffers 1-5 (SCXx1-SCXx5) were all composed of 20% (v/v) acetonitrile and 0.5% (v/v) formic acid (186006691, Waters™ Corp), with increasing concentrations of ammonium acetate (10365260, Thermo Fisher Scientific) (SCXx1, 75 mM; SCXx2, 125 mM; SCXx3, 200 mM; SCXx4, 300 mM; and SCXx5, 400 mM). Buffer 6 (Buffer X) was composed of 5% (v/v) ammonium hydroxide (221228-1L-M, Merck) and 80% (v/v) acetonitrile. 60 µl SCXx1 was first added to each stage tip. Stage tips were then centrifuged, and elution fraction 1 was collected. Each stage tip was subsequently transferred to a fresh microcentrifuge tube with a small hole cut in the lid, and the process was repeated for buffers SCXx2-SCXx5 and Buffer X to elute fractions 2-6. For each sample, elution fractions were then combined into 3 pools (pool 1, fractions 1 and 2; pool 2, fractions 3 and 4; and pool 3, fractions 5 and 6). The pooled fractions were then incubated at 45 °C in a vacuum concentrator until all liquid was completely evaporated, and were subsequently stored at -80 °C.

##### 3.19.4 LC-MS/MS analysis

Peptides were analysed by liquid chromatography with mass spectrometry using Q Exactive coupled to an Easy-nLC 1000 HPLC (ThermoFisher Scientific) equipped with a 100-µm internal dimension × 2-cm Acclaim PepMap 100 Precolumn (ThermoFisher Scientific) and a 75-µm internal dimension × 50-cm, 2-µm particle EASY-Spray HPLC C18 analytical column (ThermoFisher Scientific). Analytical solvents A (0.1% FA) and B (80% acetonitrile plus 0.1% FA). 1µg peptides were separated at a 250 nl/min flow rate by a 180 min analytical gradient which was 5%–35% solvent B over 161 minutes, rising to 99% solvent B over 1 minutes, followed by a 14 minute wash at 99% solvent B. Analytical column was held at 45°C. Data were acquired in data-dependent acquisition mode with the following settings: MS1 acquisition settings: 350–1800 m/z window, 70,000 resolution, 1x10e6 automatic gain control (AGC) target, 20 ms maximum injection time; MS2 acquisition settings: 35,000 resolution, quadrupole isolation at an isolation width of m/z 1.6, high-energy collisional dissociation fragmentation (NCE 31) with fragment ions scanning in the Orbitrap from m/z 110, 2x10e5 AGC target, 120 ms maximum injection time. Dynamic exclusion was set to ±10 ppm for 15 seconds. MS2 fragmentation was triggered on precursors 2×10e2 counts and above.

##### 3.19.5 Data processing and analysis

Raw files were analysed with MaxQuant (v1.6.1.0) with integrated Andromeda used for searching MS spectra. The following parameters were used: specific trypsin digestion; up to 2 missed cleavages allowed; Precursor ion tolerance: 20 and 4.5 ppm for first and main searches, respectively; Human database (UniProt SwissProt Nov 2018) with additional inbuilt MaxQuant contaminant database (containing 246 common contaminants); oxidation (M) and N terminal acetylation as variable modifications; carbamidomethylation (C) as a fixed modification; Match between runs enabled. Proteins were filtered for FDR (< 1%). Further analyses were carried out in Perseus (v1.6.0.2):^71^ proteins identified by site, common contaminants and reverse sequences were removed; log 2 transformation on all values followed by row-wise mean and column-wise median normalisation before filtering for 8 valid values across all conditions (total 8 conditions); volcano plot cut offs were set as fdr = 0.015 and s0 = 1 for 4.5 hours infection versus no infection, and fdr = 0.012 and s0 = 1 for 6 hours infection versus no infection.

#### 3.20 Molecular biology

##### 3.20.1 Streaking of bacterial stabs

Bacteria in stab culture transformed with pcDNA-3xFLAG-Pontin (51635, Addgene) or pcDNA-3xHA-Reptin (51636, Addgene) were struck onto LB agar plates containing 100 µg/ml ampicillin (A9518-25G, Merck) into single colonies. The plates containing the transformed bacteria were incubated (agar side up) overnight at 37 °C.

##### 3.20.2 Transformation of plasmid DNA into competent cells

Plasmid DNA was transformed into TOP10 chemically competent E.coli (C404003, Thermo Fisher Scientific). For each transformation, 50 ng of plasmid DNA was mixed with 50 µl competent cells and subsequently incubated on ice for 20 minutes. Cells were then heat-shocked by incubating for 45 seconds at 42 °C while shaking at 300 rpm. Cells were then incubated on ice for 6 minutes, and subsequently topped up to a total volume of 500 µl with pre-warmed LB broth (51208-500G-F, Merck) (without antibiotic). The cell suspension was then incubated for 1 hour at 37 °C while shaking at 300 rpm, plated out onto LB agar (19344-500G-F, Merck) containing 100 µg/ml ampicillin, and subsequently incubated (agar side up) overnight at 37 °C. The same protocol was also performed for the transformation of ligation products from side-directed mutagenesis (section 3.20.9), except 25 ng (5 µl) plasmid DNA was transformed rather than 50 ng.

##### 3.20.3 Production of glycerol stocks

To produce glycerol stocks, a single colony of the bacteria transformed with the desired plasmid was used to inoculate 1 ml LB broth containing 100 µg/ml ampicillin. This culture was incubated for 8 hours at 37 °C while shaking at 200 rpm, and subsequently topped up to a total volume of 7 ml with pre-warmed LB broth containing 100 µg/ml ampicillin. This culture was then incubated overnight at 37 °C while shaking at 200 rpm. The following day, 0.5 ml sterile 50% (v/v) glycerol was added to a cryovial tube. 0.5 ml bacterial culture was then added to the cryovial tube containing 0.5 ml glycerol, vortexed, and immediately stored at - 80 °C.

##### 3.20.4 Small scale plasmid DNA preparation

To purify plasmid DNA on a small scale, a single colony of the bacteria transformed with the desired plasmid was used to prepare a 7 ml overnight culture as described above. Following the overnight incubation and potential glycerol stock preparation, 5 ml of the bacterial culture was then centrifuged at 2,300 x g for 15 minutes at RT, and the supernatant was removed and discarded. The culture pellet was subsequently processed for plasmid DNA purification using the Monarch® Plasmid DNA Miniprep Kit (T1010L, New England Biolabs) according to the manufacturer’s instructions, except plasmid DNA was eluted in distilled water (the kit elution buffer contains EDTA, which is incompatible with subsequent sequencing techniques).

##### 3.20.5 Large scale plasmid DNA preparation

To purify plasmid DNA on a large scale, a sterile loop was first used to take a small sample (∼20 µl) from the glycerol stock containing the bacteria transformed with the desired plasmid. This sample was then used to inoculate 2 ml LB broth containing 100 µg/ml ampicillin. This culture was incubated for 8 hours at 37 °C while shaking at 200 rpm, and subsequently transferred to 150 ml LB broth containing 100 µg/ml ampicillin. This culture was then incubated overnight at 37 °C while shaking at 200 rpm. Following the overnight incubation, the 150 ml bacterial culture was then centrifuged at 2,300 x g for 15 minutes at RT. The supernatant was removed and discarded, and the culture pellet was subsequently processed for plasmid DNA purification using the PureLink™ HiPure Plasmid Maxiprep Kit (K210007, Thermo Fisher Scientific) according to the manufacturer’s instructions.

##### 3.20.6 Quantification of plasmid DNA

All purified plasmid DNA samples were quantified using the dsDNA setting on a NanoDrop™ One microvolume UV-vis spectrophotometer. For accuracy, 3 readings were taken for each sample, which were subsequently averaged to obtain the average DNA concentration (ng/µl).

##### 3.20.7 DNA agarose gel electrophoresis

DNA gel electrophoresis was performed using 1% (w/v) agarose gels (1 g agarose in 100 ml tris-acetate-EDTA (TAE; BP1332-1, Thermo Fisher Scientific) buffer, with 5 µl Midori Green (S6-0016, Geneflow) to visualise DNA). DNA was prepared using 6X purple loading dye without SDS (B7025S, New England Biolabs). Quick-Load® 1kb Plus DNA Ladder (N0469S, New England Biolabs) was used for size markers. Gels were electrophoresed at 100 V for 45-60 minutes, and DNA bands were subsequently revealed using a Syngene™ G:BOX Chemi XRQ image analyser.

##### 3.20.8 Gel purification of DNA

DNA fragments were purified from 1% (w/v) agarose gels using the Monarch® DNA Gel Extraction Kit (T1020L, New England Biolabs) according to manufacturer’s instructions. DNA fragments were eluted in 10 µl RNase free water.

##### 3.20.9 Site-directed mutagenesis

Inverse PCRs for site-directed mutagenesis were assembled at room temperature using the Novagen® KOD Hot Start DNA Polymerase Kit (71086-4, Merck). For each reaction, the following components were assembled: 5 µl 10X buffer for KOD Hot Start DNA polymerase (final concentration 1X), 3 µl 25 mM magnesium sulfate (final concentration 1.5 mM), 5 µl 2 mM each dNTPs (final concentration 0.2 mM each), 1.5 µl 10 µM forward primer, 1.5 µl 10 µM reverse primer (final concentration 0.3 µM each), 1 µl 100 ng/µl template DNA (final concentration 2 ng/µl), 1 µl KOD Hot Start DNA polymerase (1 U/µl) (final concentration 0.02 U/µl) and RNase free water to a total volume of 50 µl. Details of primers used are shown in the Key Resources Table. Reactions were cycled on an Applied Biosystems™ ProFlex PCR System (4484073, Thermo Fisher Scientific) using the protocol described below.

| Step | Temperature (°C) | Time (minutes) | Number of cycles |
| --- | --- | --- | --- |
| 1. Polymerase activation | 95 | 02:00 | X1 |
| 2. Denaturation | 95 | 00:20 | X40 |
| 3. Annealing | Annealing Temp | 00:10 |  |
| 4. Extension | 70 | 00:25 per 1 kb |  |
| 5. Final extension | 70 | 10:00 | X1 |
| 6. Hold | 4 | Hold | X1 |

To digest methylated plasmid template DNA, 0.5 µl DpnI (R0176, New England Biolabs) was added to each 50 µl reaction, mixed, and incubated for 4 h at 37 °C. Samples were then purified using the Monarch® PCR & DNA Cleanup Kit according to manufacturer’s instructions. PCR products were eluted in 40 µl RNase free water.

Reaction mixes for 5’ end phosphorylation of PCR products were assembled at room temperature. For each reaction, the following components were assembled: 39 µl eluted PCR product, 5 µl 10X T4 polynucleotide kinase reaction buffer (B0201S, New England Biolabs; final concentration 1X), 0.5 µl 100 mM ATP (366-10001-1, RayBiotech; final concentration 1 mM), and 1 µl T4 polynucleotide kinase (M0201S, New England Biolabs) to have 50 µl in total. Reactions were incubated for 1 h at 37 °C, and subsequently purified using the Monarch® PCR & DNA Cleanup Kit according to manufacturer’s instructions. PCR products were eluted in 10 µl RNase free water, and were quantified using the dsDNA setting on a NanoDrop™ One microvolume UV-vis spectrophotometer.

Reaction mixes for self-ligation of plasmid DNA were prepared using the Quick Ligation™ Kit (M2200L, New England Biolabs) according to manufacturer’s instructions. For each reaction, the following components were assembled: 50 ng eluted PCR product, 10 µl 2X Quick Ligation™ buffer (final concentration 1X), 1 µl Quick T4 DNA ligase, and RNase free water to a total volume of 20 µl. Reactions were incubated for 5 minutes at 25 °C, chilled on ice, and subsequently transformed into TOP10 competent *E.coli* as described in section 3.20.2. Following transformation, liquid cultures and glycerol stocks were made (section 3.20.3), and plasmids were purified (section 3.20.4) and sent for sequencing as described in section 3.20.11. Maxipreps of sequence-verified clones were then prepared as described in section 3.20.5.

To generate the replication-defective RV-A16 nanoluciferase replicon, plasmid DNA encoding an RV-A16 nanoluciferase replicon (pRVA-16-NL), in which the P1 region is replaced by the NanoLuc® luciferase reporter gene under the control of the RV-A16 IRES (gift from Stanley Lemon and Kevin McKnight, University of North Carolina), was used as a template.^68^ The conserved 3D polymerase active-site motif YGDD was mutated to YGAA by site-directed mutagenesis as described above, and constructs were sequence-verified across the 3D region (Eurofins) as described in section 3.20.11. The plasmid encoding the replication-defective replicon (hereby referred to as pRVA-16-NL-YGAA) was subsequently linearised, and mutant RNA was *in vitro* transcribed from the linearised plasmid and purified as described in section 3.20.12.

To generate single, double and quadruple RV-A16 mutants (section 3.21.4), plasmid DNA encoding the full-length wild-type RV-A16 genome (pR16.11) was first used as a template for the generation of single mutants (VP1 V285A, 2A Y92H, 2C M121V, and 2C T284S) by site-directed mutagenesis as described above. Subsequently, pRV-A16 2A Y92H was used as a template for the generation of pRV-A16 VP1-2A DM, and pRV-A16 2C T284S was used as a template for the generation of pRV-A16 2C-DM (double mutants), by site-directed mutagenesis as described above. To generate the quadruple mutant containing all 4 mutations, pRV-A16 VP1-2A DM and pRV-A16 2C-DM were each digested using ApaI (R0114S, New England Biolabs) and ClaI (R0197S, New England Biolabs) (1 µg plasmid DNA, 5 µl 10X rCutSmart® buffer (B6004S, New England Biolabs) (final concentration 1X), 2 µl ApaI, 2 µl ClaI, and RNase free water to a total volume of 50 µl) for 2 h at 37 °C. Digested DNA samples were then run on a 1% (w/v) agarose gel, and the large fragment for pRV-A16 VP1-2A DM (at around 10 kb, containing VP1 V285A and 2A Y92H) and the small fragment for RV-A16 2C-DM (at around 1.2 kb, containing 2C M121V and 2C T284S) were both excised and gel-purified as described in section 3.20.8. Fragments were then ligated using the Quick Ligation™ Kit according to manufacturer’s instructions. Reactions were assembled as follows: 10 µl Quick Ligation™ buffer (final concentration 1X), 5 µl pRV-A16 VP1-2A DM fragment (the “vector”), 5 µl RV-A16 2C-DM fragment (the “insert”), and 1 µl Quick T4 DNA ligase. A vector-only reaction control, in which the 2C-DM fragment was replaced with RNase free water, was also prepared. Reactions were incubated for 5 minutes at 25 °C, chilled on ice, and subsequently transformed into TOP10 competent *E.coli* as described in section 3.20.2. Plasmids were sequenced (Eurofins) at each step (single mutants, double mutants, quadruple mutant) to confirm substitution of the required nucleotide at the correct position, as described in section 3.20.11. The plasmids encoding the site-directed mutants were subsequently linearised, and mutant viral RNA was *in vitro* transcribed from the linearised plasmids and purified as described in section 3.20.12. Mutant viral RNA was then transfected into HeLa-H1 cells to generate virus stocks as described in section 3.11.

##### 3.20.10 In vivo DNA assembly

The plasmid encoding Myc-GFP-2BC3ABCD (pRK5-Myc-GFP-2BC3ABCD) was generated by *in vivo* DNA assembly^82^ as follows. Plasmid DNA encoding the full-length wild-type RV-A16 genome (pR16.11), and plasmid DNA encoding Myc-GFP (pICC1564), were used as templates. Single-tube PCRs (in which all primers and vectors were added into a single reaction) were assembled at room temperature using the Novagen® KOD Hot Start DNA Polymerase Kit as described in 3.20.9, with minor modifications. For each reaction, the following components were assembled: 2.5 µl 10X buffer for KOD Hot Start DNA polymerase (final concentration 1X), 2 µl 25 mM magnesium sulfate (final concentration 2 mM), 2.5 µl 2 mM each dNTPs (final concentration 0.2 mM each), 0.5 µl DMSO (final concentration 2%, v/v), 1 µl 5 µM pR16.11 forward primer, 1 µl 5 µM pR16.11 reverse primer, 1 µl 5 µM pICC1564 forward primer, 1 µl 5 µM pICC1564 reverse primer (final concentration 0.2 µM each), 2.5 µl 1 ng/µl pR16.11 template DNA, 2.5 µl 1 ng/µl pICC1564 template DNA (final concentration 0.1 ng/µl each), 0.5 µl KOD Hot Start DNA polymerase (1 U/µl) (final concentration 0.02 U/µl) and RNase free water to a total volume of 25 µl. Details of primers used are shown in the Key Resources Table. Reactions were cycled on an Applied Biosystems™ ProFlex PCR System as outlined in 3.20.9, using an annealing temperature of 57 °C, an extension time of 3 minutes 30 seconds, and X20 cycles of denaturation, annealing and extension instead of X40 cycles (less cycles to minimise the chance of polymerisation errors).

5 µl PCR product was then run on a 1% (w/v) agarose gel (as outlined in section 3.20.7) to confirm amplification of correctly sized fragments. To digest methylated plasmid template DNA, 1 µl DpnI was added to the remaining PCR product, mixed, and incubated for 1 h at 37 °C. 10 µl DpnI-treated PCR product was then directly transformed into TOP10 competent *E.coli* as described in section 3.20.2. Following transformation, liquid cultures were made (section 3.20.3) and verified by colony PCR. For each reaction, the following components were assembled: 0.5 µl 10 µM forward primer, 0.5 µl 10 µM reverse primer (final concentration 0.2 µM each), 12.5 µl OneTaq® Quick-Load® 2X Master Mix with Standard Buffer (M0486L, New England Biolabs), RNase-free water to a total volume 25 µl, and template DNA (from the same colony used to make the liquid culture; the tip used to pick the colony was briefly placed inside the reaction tube and turned 5 times). Details of primers used are shown in the Key Resources Table. Reactions were cycled on an Applied Biosystems™ ProFlex PCR System as outlined below

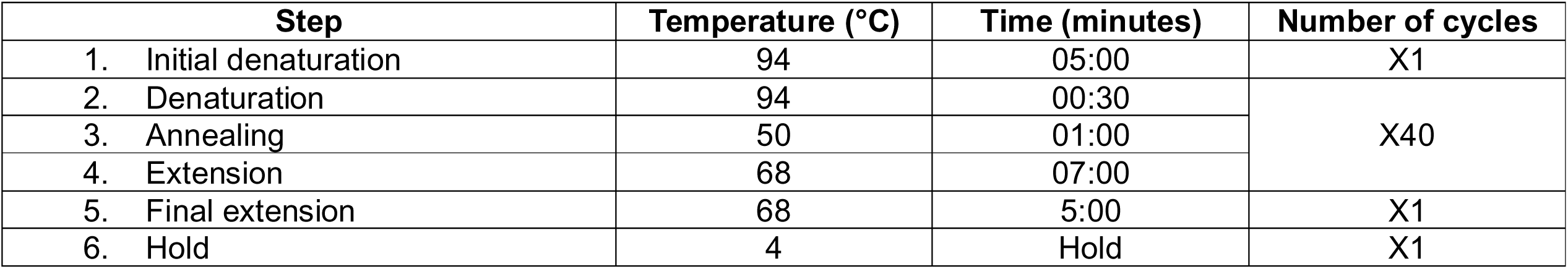

Colony PCR product was then run on a 1% (w/v) agarose gel (as outlined in section 3.20.7) to confirm amplification of correctly sized fragments. The next day, for clones verified by colony PCR, glycerol stocks were made (section 3.20.3) and plasmids were purified (section 3.20.4). Purified plasmid DNA was then verified by analytical digest as follows: 1 µg plasmid DNA, 2.5 µl 10X rCutSmart® Buffer (final concentration 1X), 0.5 µl EcoR1-HF® (R3101L, New England Biolabs), and RNase free water to a total volume of 25 µl. Reactions were incubated for 15 minutes at 37 °C (EcoR1-HF® is a time-saver enzyme), and correct digestion was confirmed by running samples on a 1% (w/v) agarose gel (as outlined in section 3.20.7). Clones verified by analytical digest were sent for sequencing (Eurofins) as described in section 3.20.11. A maxiprep of the sequence-verified clone was then prepared as described in section 3.20.5.

##### 3.20.11 Sequencing of plasmid DNA

For each sample to be sent for sequencing, 1 µg purified plasmid DNA was diluted in distilled water to have a final volume of 20 µl (50 ng/µl). Sequencing of purified plasmid DNA was performed by Eurofins.

##### 3.20.12 In-vitro transcription and RNA purification

Reaction mixes for linearisation of plasmid DNA by digestion with the indicated enzymes were assembled as follows: 10 µg plasmid DNA, 10 µl 10X rCutSmart® Buffer (final concentration 1X), 2 µl enzyme, and RNase free water to a total volume of 100 µl. The plasmid encoding HeLa-adapted RV-C15 (pC15-Rz-K41) was digested with BstB1 (R0519S, New England Biolabs). The plasmids encoding HeLa-adapted RV-C41 (pC41-Rz-K41), full-length RV-A16 (pR16.11) and the pRV-A16 mutant plasmids (VP1 V285A, 2A Y92H, 2C M121V, 2C T284S, VP1-2A DM, 2C-DM, and the quadruple mutant) were digested with SacI-HF® (R3156S, New England Biolabs). The plasmids encoding full-length RV-A1a (pWIN1a) and full-length RV-B14 (pWR3.26) were digested with MluI-HF® (R3198S, New England Biolabs). The plasmid encoding the replication-defective RV-A16 nanoluciferase replicon (pRVA-16-NL-YGAA) was digested with BamH1-HF® (R3136S, New England Biolabs). Reactions were incubated for 2 hours at 37 °C and, if necessary, were subsequently heat-inactivated for 20 minutes at 65 °C (only required for SacI-HF®).

Linearised plasmid DNA was then purified using the QIAquick PCR Purification Kit (28106, Qiagen) according to manufacturer’s instructions. Purified DNA was eluted in 20 µl RNase free water, and was quantified using the dsDNA setting on a NanoDrop™ One microvolume UV-vis spectrophotometer. Successful linearisation was confirmed by running samples on a 1% (w/v) agarose gel (as described in section 3.20.7) and comparing digested plasmid DNA to an undigested control.

*In vitro* transcription was performed using the MEGAscript™ T7 Transcription Kit (AMB13345, Thermo Fisher Scientific) according to manufacturer’s instructions. Reactions were assembled at room temperature as follows: 1 µg purified linearised plasmid DNA template, 2 µl ATP, 2 µl CTP, 2 µl GTP, 2 µl UTP, 2 µl 10X reaction buffer, 2 µl RNA polymerase, and RNase free water to a total volume of 20 µl. Reactions were incubated for 4 h at 37 °C. Subsequently, 1 µl TURBO DNase was added to each reaction, mixed well, and incubated for 15 minutes at 37 °C. *In vitro* transcribed RNA was then purified using magnetic SPRI RNA purification beads, which were made in-house according to established protocols.^61^ Beads were resuspended by gentle inversion, and a volume of beads equivalent to the volume of sample was added to each sample and incubated for 5 minutes at room temperature. Samples were placed on a magnetic rack until the beads migrated to the holder, and the clear supernatant was removed. Beads (with RNA bound) were then washed 2X in 35 µl 70% (v/v) ethanol, and RNA was subsequently eluted in 20 µl RNase free water. RNA was quantified using the RNA setting on a NanoDrop™ One microvolume UV-vis spectrophotometer. Purified RNA was then transfected into HeLa cells either directly for experiments (sections 3.9 and 3.10) or to generate virus stocks (section 3.11).

#### 3.21 RV passaging with CB-6644 and mutant analysis

##### 3.21.1 Passaging of RV in drug-treated cells

HeLa-H1 cells were seeded in complete medium into 25 cm^2^ cell culture flasks. At 95-100% confluence, cells were infected with RV-A16 (MOI=0.1, diluted in infection medium) alone or in the presence of 15 nM CB-6644 or 92 nM NITD008, and incubated at 33 °C, 5% CO_2_ until 100% of cells exhibited CPE. After three cycles of freezing and thawing to release viral particles, cell lysates were clarified by centrifugation (2,300 x g for 30 minutes at 4 °C) and supernatants were collected (passage 1 virus; P1 virus). Fresh HeLa-H1 cells seeded in 25 cm^2^ cell culture flasks were then infected at 95-100% confluence with P1 viruses (MOI=0.1, assuming a titre of 1.0x10^7^) alone or in the presence of 15 nM CB-6644 or 92 nM NITD008, and incubated at 33 °C, 5% CO_2_ until 100% of cells exhibited CPE. After three cycles of freezing and thawing, cell lysates were clarified by centrifugation as above and supernatants were collected (P2 virus). For CB-6644-passaged RV-A16, this process was repeated up to P22, with CB-6644 concentrations being increased stepwise (30 nM at P3, 60 nM at P16, and 100 nM at P19). For NITD008-passaged RV-A16, this process was repeated up to P15, with NITD008 concentrations being increased stepwise (200 nM at P11, 400 nM at P14). For untreated-passaged RV-A16, this process was repeated up to P22. For each virus, the final passage (P22 or P15) was performed in a T175 flask to maximise virus yield. After three cycles of freezing and thawing, cell lysates were clarified by centrifugation as described in section 2.3.1, and supernatants were aliquoted and stored at -80 °C. For each virus (CB-6644 P22, NITD008 P15, and Untreated P22), 1 aliquot was kept for analysis for virus titre by TCID_50_ assay.

##### 3.21.2 Dose effect assay and extraction of RNA from virions

To assess whether passaged viruses had evolved resistance to their respective drugs, passaged viruses were challenged in a CB-6644 dose effect assay (Figure 4C, CB-6644 P22 and Untreated P22) or in a NITD008 dose effect assay (Figure S2F, NITD008 P15 and Untreated P22) as described in section 3.4. For the CB-6644 P22 virus, the IC_90_ and IC_99_ was calculated and compared to that of the Untreated P22 virus.

To identify mutations giving rise to CB-6644 resistance, the CB-6644 P22 virus was selected for sequencing. Unpassaged RV-A16 (the parental stock used to generate CB-6644 P22) was also selected for sequencing. Viral RNA was extracted from virions as follows. 1 ml virus stock was added to 3 ml TRI Reagent® LS (T3934, Merck), vortexed for 10 seconds, and incubated for 2 minutes at room temperature. 800 µl chloroform (32211-M, Merck) was then added, and the mixture was vortexed for 10 seconds until cloudy pink and then centrifuged (2,300 x g for 10 minutes at room temperature). The 1,800 µl aqueous phase was then transferred to 2X 2 ml screw cap tubes (900 µl per tube), taking care not to disturb the white layer containing DNA. 900 µl isopropanol (34863, Merck) and 1 µl Glycoblue™ (AM9515, Thermo Fisher Scientific) was then added to each tube, and the mixtures were vortexed for 10 seconds and incubated for 1 hour at -80 °C to precipitate RNA. Tubes were subsequently centrifuged (15,000 x g for 15 minutes at 4 °C). The isopropanol was then carefully removed, and the pellets were washed in 175 µl 70% (v/v) ethanol and centrifuged (15,000 x g for 3 minutes at 4 °C). The 70% ethanol was then carefully removed, and 1 of the 2 pellets was resuspended in 20 µl RNase free water. This resuspended RNA was then transferred to the other tube and used to resuspend the second pellet. RNA for each sample was quantified using the RNA setting on a NanoDrop™ One microvolume UV-vis spectrophotometer.

##### 3.21.3 Sequencing of viral RNA

Virion RNA sequencing libraries were prepared using a cDNA-PCR method and sequenced on a P2 Solo instrument (Oxford Nanopore Technologies). Briefly, 500 ng of purified RNA was reverse transcribed using an oligo(dT)-anchored primer and a strand-switching strategy to preferentially generate full-length cDNA transcripts from polyadenylated RNA molecules. Second-strand synthesis and PCR amplification were then performed according to the manufacturer’s protocol (cDNA-PCR Sequencing Kit, SQK-PCB114-24). Following amplification, a post-PCR cleanup step using AMPure XP beads at a 1:1 bead-to-sample ratio was applied to reduce short artefactual fragments prior to the ligation of sequencing adapters. The final library was loaded onto a P2 flow cell (FLO-PRO114M) incorporating R10.4.1 pore chemistry and sequenced for 1 hour using MinKNOW (v24.06.16) with default acquisition parameters, generating a total of 1.95 million reads. Raw POD5 files were converted to FASTQ format using Dorado basecaller (v0.7.1) with a high accuracy model. Basecalled reads were aligned to a custom reference genome using Bowtie2 (v2.4.4)^70^ with default parameters. The reference was constructed from the wild-type stock virus sequence, which served as the canonical ensemble for all downstream analyses. Mutant viral RNA sequences were aligned to this reference to assess sequence variation and transcriptomic differences. Alignment quality and coverage were evaluated using SAMtools (v1.13),^72^ and alignments were visualised using the Integrated Genome Viewer (IGV). An allele frequency threshold of 0.5 was applied in order to identify nucleotide substitutions occurring in at least >50% of reads at each position. 4 missense mutations were identified, giving rise to the following 4 amino acid changes: VP1 V285A, 2A Y92H, 2C M121V, and 2C T284S.

##### 3.21.4 Generation of RV mutants

Based on the sequencing data, RV-A16 single mutants (VP1 V285A, 2A Y92H, 2C M121V, and 2C T284S), double mutants (VP1-2A DM and 2C-DM) and a quadruple mutant (containing all 4 mutations) were generated by site-directed mutagenesis as described in section 3.20.9. Mutant viral RNA was *in vitro* transcribed and purified as described in section 3.20.12, and transfected into HeLa-H1 cells to generate virus stocks as described in section 3.11. The mutant challenge experiment (Figure 4D) was performed as described in section 3.3.

### 4 Quantification and statistical analysis

#### 4.1 Statistical analysis: Prism

For all data except Figure 1B-C, data was analysed using the appropriate statistical test (two-tailed paired *t*-test, one-way ANOVA, two-way ANOVA) and *post-hoc* test (Dunnett’s, Holm-Sidak’s, Tukey’s) as outlined in the figure legend and using Prism software (v10.5.0) (GraphPad). *, *P* < 0.05; **, *P* < 0.01; ***, *P* < 0.001; ****, *P* < 0.001; ns, not significant.

##### 4.1.1 IC_50_, IC_90,_ and IC_99_ calculation

For CB-6644 (Figure 1F, Figure 4C) dose effects, the IC_50_, IC_90_ and IC_99_ values were determined using Prism software (v10.5.0) (GraphPad), in XY table format. For each experiment, RV-A16 titres (TCID_50_/ml) at each drug concentration were normalised to the titre in the DMSO-treated control. Normalised titres and corresponding drug concentrations were arranged in descending order (starting from the highest concentration). Drug concentrations were log-transformed [X=log(X)].

IC_50_ values (Figure 1F), were calculated using the *log(inhibitor) vs. normalized response - variable slope (4 parameters)* function. IC_90_ and IC_99_ values (Figure 4C) were calculated using the *log(agonist) vs. response - Find ECanything* function, with the constraint parameter *F* (equal to) set to 10 for IC_90_ and 1 for IC_99_.

#### 4.2 Statistical analysis: proteomics data

Proteomics data (Figure 1B-C) was analysed as outlined in section 3.19.5.

## Notes

### Competing Interest Statement

The authors have declared no competing interest.

