## Supplementary Figures for "A host ATPase essential for rhinovirus replication is an antiviral target with a high barrier to resistance"

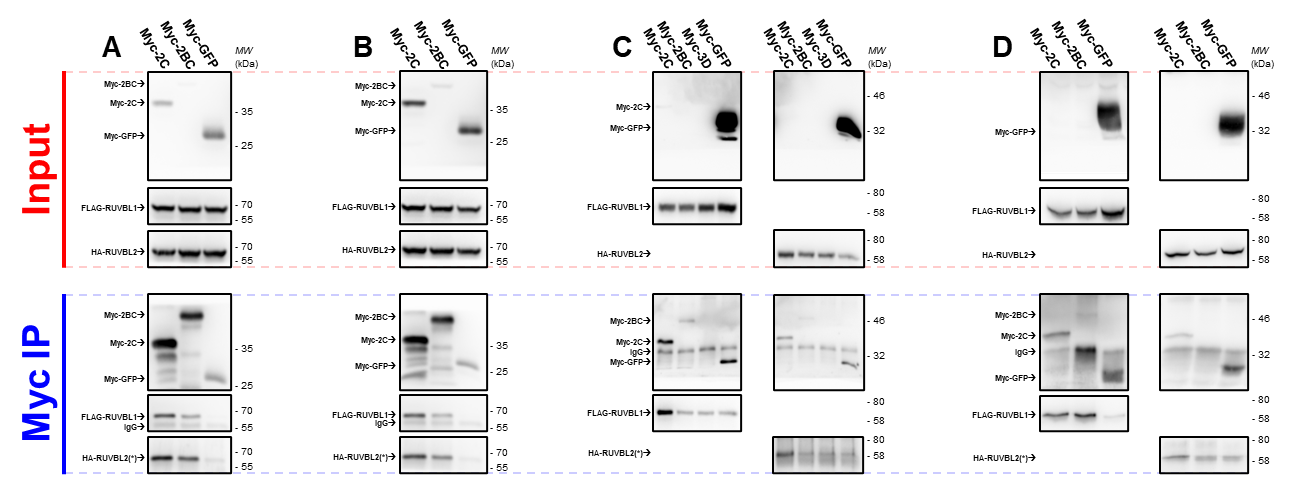

**Supplementary Figure 1, related to figure 1H.**

**(A-D) RUVBL1 and RUVBL2 co-immunoprecipitate with RV-A16 2C or 2BC in the absence of other viral components, related to figure 1H (showing the independent experimental replicates).** HeLa-H1 cells were transfected with constructs encoding Myc-tagged RV NSPs (2C or 2BC) or Myc-GFP with both FLAG-RUVBL1 and HA-RUVBL2 (triple transfection, **A-B**), or with either FLAG-RUVBL1 or HA-RUVBL2 (double transfection, **C-D**). After 30 h (**A-B**) or 48 h (**C-D**), Myc-tagged proteins were immunoprecipitated from cell lysates. Cell lysates (input) and immunoprecipitated fractions (α-Myc IP) were analysed by western blotting for Myc, FLAG, and HA. (*) HA-RUVBL2 overlaps with IgG heavy chain. FLAG-RUVBL1 and HA-RUVBL2 with Myc-2C, N=4 (**A-D**); FLAG-RUVBL1 with Myc-2BC, N=3 (**A-B, D**); HA-RUVBL2 with Myc-2BC, N=2 (**A-B**).

N=number of independent experiments.

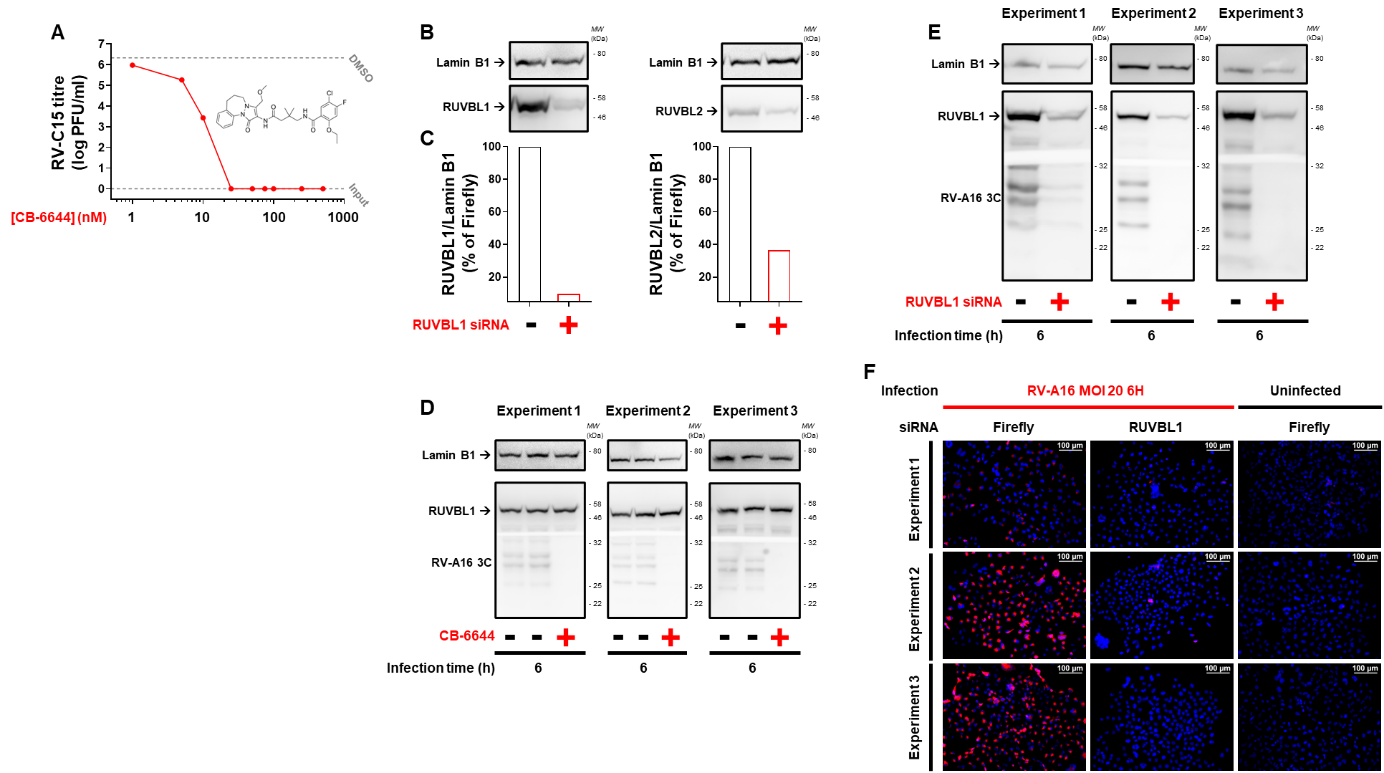

**Supplementary Figure 2, related to figures 2 and 3.**

**(A) CB-6644 inhibits RV-C15 replication, related to figure 2C.** HeLa-E8 cells were infected with RV-C15 (MOI 1) and treated at 1 h post-infection with DMSO or increasing concentrations of CB-6644 (1, 5, 10, 25, 50, 75, 100, 250 and 500 nM). Viral titres were quantified at 0 h and 16 h (N=1). Viral titres in CB-6644-treated cells at 16 h are shown as individual points with means connected by a line. Mean viral titres of untreated cells at 0 h (input) and of DMSO-treated cells at 16 h are represented by dashed lines.

**(B-C) siRNA knockdown of RUVBL1 reduces expression of both RUVBL1 and RUVBL2, related to figure 3F. (B)** HeLa-H1 cells were transfected with 10 nM siRNA targeting RUVBL1 or firefly luciferase. At 72 h, lysates were analysed by western blotting for RUVBL1, RUVBL2 and lamin-B1 (N=1). **(C)** RUVBL1 and RUVBL2 expression was quantified and normalised to lamin-B1.

**(D) CB-6644 inhibits RV NSP production, related to figure 3B (showing the 3 independent experiments).** HeLa-H1 cells infected with RV-A16 (MOI 20) were treated at 1 h post-infection with DMSO or 500 nM CB-6644. At 6 h, lysates were analysed by western blotting for RV-A16 3C, RUVBL1, and lamin-B1.

**(E) siRNA knockdown of RUVBL1 inhibits RV NSP production, related to figure 3F (showing the 3 independent experiments).** HeLa-H1 cells were transfected with 10 nM siRNA targeting RUVBL1 or firefly luciferase for 72 h and then infected with RV-A16 (MOI 20). At 6 h, lysates were analysed by western blotting for RV-A16 3C, RUVBL1, and lamin-B1.

**(F) siRNA knockdown of RUVBL1 inhibits RV NSP production, related to figure 3I (showing the 3 independent experiments).** HeLa-H1 cells were transfected with 10 nM siRNA targeting RUVBL1 or firefly luciferase for 72 h and then infected with RV-A16 (MOI 20) or left uninfected. At 6h, cells were stained for RV-A16 2C (red) and nuclei (DAPI, blue). Representative images are shown.

N=number of independent experiments.

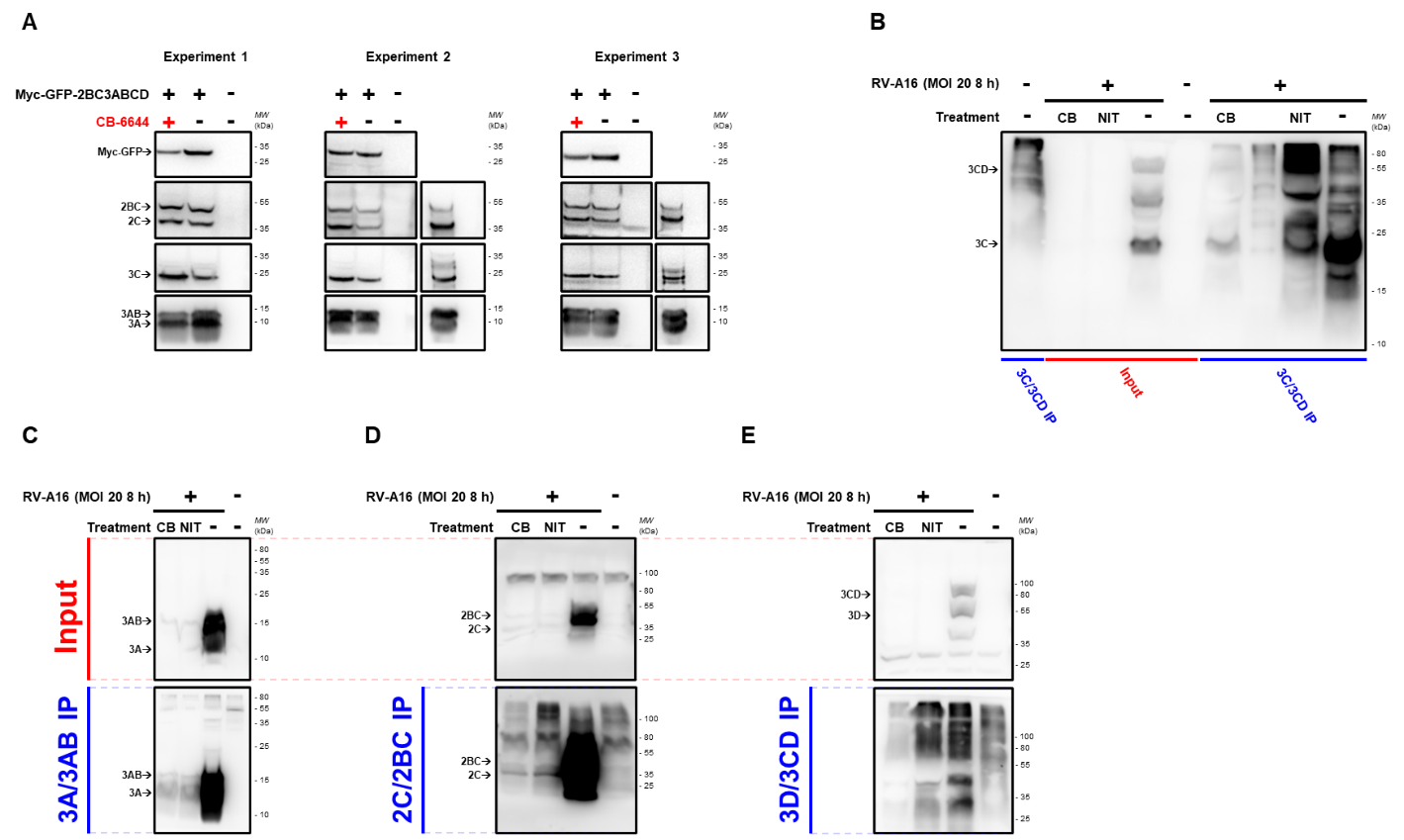

**Supplementary Figure 3, related to figure 3O.**

**(A) RUVBL1/2 is not required for the cleavage of a transfected RV-A16 polyprotein fragment, related to figure 3O (showing the 3 independent experiments).** HeLa-H1 cells were transfected with a construct encoding Myc-GFP-2BC3ABCD in the presence of DMSO or 500 nM CB-6644 for 21 h or were left untransfected (N=3). In parallel, HeLa-H1 cells were infected with RV-A16 for 8 h or were left uninfected (N=2). At 21 and 8 h, respectively, lysates were analysed by western blotting for Myc-GFP and RV-A16 2C, 3A and 3C.

**(B-E) RV-A16 polyprotein cleavage in infected cells treated with CB-6644.** HeLa-H1 cells were infected with RV-A16 (MOI 20) and treated after virus adsorption with 500 nM CB-6644 (CB) or 5 µM NITD008 (NIT), which inhibits viral RNA replication. Alternatively, cells were infected without any treatment or left uninfected. After 8 h, the indicated RV NSPs were immunoprecipitated from cell lysates. Cell lysates (input) and immunoprecipitated fractions (IP) were analysed by western blotting for the indicated RV NSPs (N=1 each for 3A/3AB, 3C/3CD, 2C/2BC, and 3D/3CD).

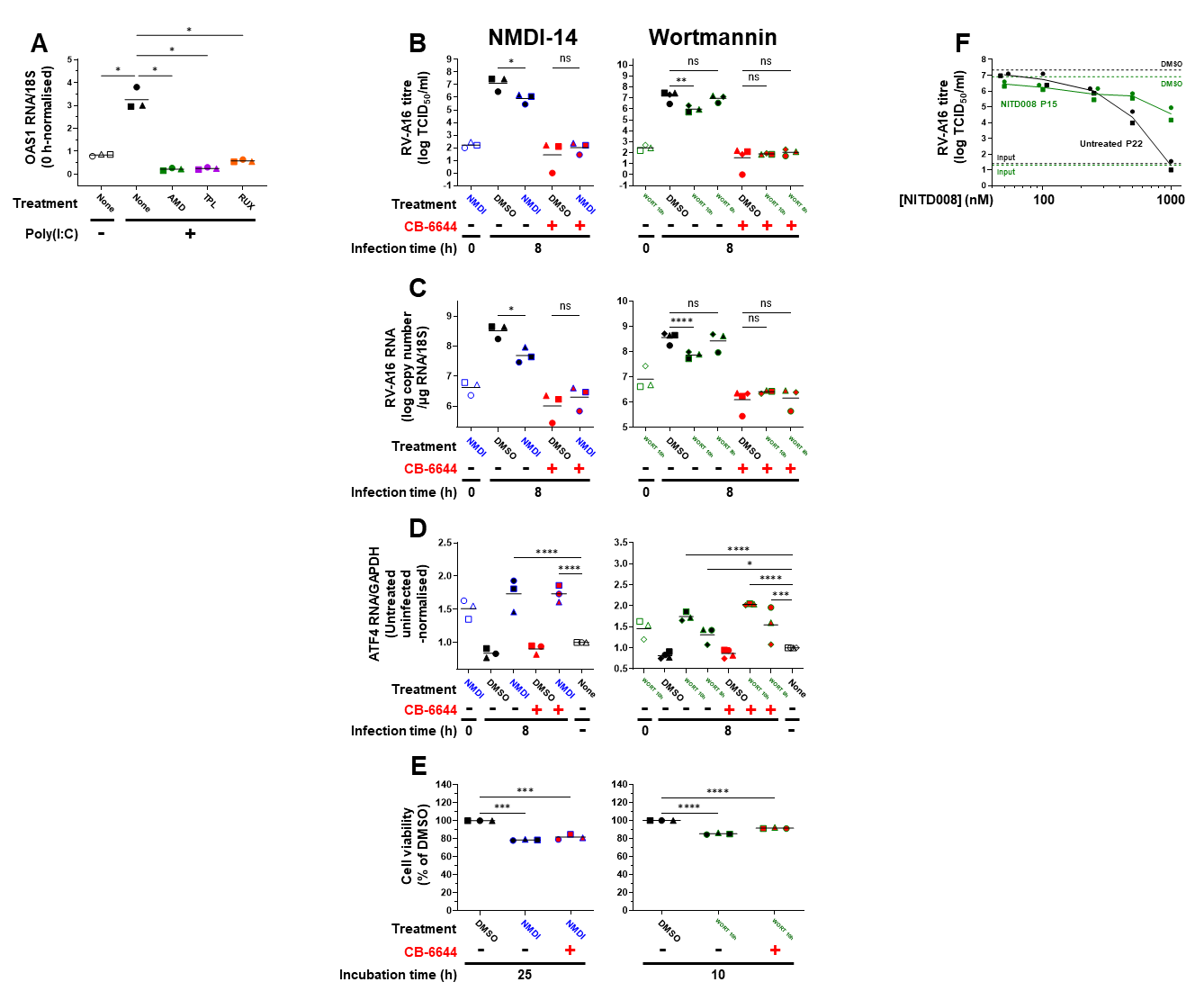

**Supplementary Figure 4.**

**(A) Inhibitors of cellular transcription are functional in HeLa-H1 cells, related to figure 4A.** HeLa-H1 cells were treated with 2 µg/ml actinomycin D (AMD), 10 µM triptolide (TPL), 10 µM ruxolitinib (RUX) or left untreated for 1 h, and subsequently treated with poly(I:C) (50 µg/ml) alone or in combination with each drug, or left untreated, for 9 h. At 10 h post-initial treatment, OAS1 mRNA levels were quantified by RT-qPCR and normalised to untreated cells at 0 h (0 h points not shown) (N=3).

**(B-E) Inhibitors of the nonsense-mediated mRNA decay pathway do not abrogate the antiviral activity of CB-6644.** HeLa-H1 cells were pre-treated with 50 µM NMDI-14 (NMDI, blue) or DMSO for 16 hours, or with 10 µM Wortmannin (WORT 10h, green) or DMSO for 1 hour, or were left untreated. Cells were subsequently infected with RV-A16 (MOI 20) alone or in the presence of the corresponding pre-treatment. Immediately after the end of the virus adsorption, cells were re-treated with the corresponding pre-treatment, or received their first treatment of 10 µM Wortmannin (WORT 8h, green) or DMSO. All treatments at this point were given in combination with 500 nM CB-6644 or DMSO. **(B)** Viral titres at 0 h and 8 h were quantified (N=3-4). **(C)** Viral RNA at 0 h and 8 h was quantified by RT-qPCR (N=3-4). **(D)** ATF4 mRNA levels at 0 h and 8 h were quantified by RT-qPCR, and were normalised to untreated uninfected cells (N=3-4). **(E)** In parallel, HeLa-H1 cells were treated as above with NMDI-14, Wortmannin, or DMSO in combination with CB-6644 or DMSO at the corresponding time. Cell viability was measured by resazurin assay at 25 h (NMDI-14) or 10 h (Wortmannin) (N=3).

**(F) NITD008-passaged RV-A16 exhibits partial resistance to NITD008, related to figure 4C.** NITD008-passaged RV-A16 was generated similarly to figure 4B, with NITD008 concentrations being increased stepwise (92 nm at passages 1-10; 200 nM at passages 11-13; 400 nM at passages 14-15). Subsequently, HeLa-H1 cells were infected with NITD008-passaged RV-A16 (P15) or untreated-passaged RV-A16 (P22, from figure 4B) (MOI 1) and treated with DMSO or increasing concentrations of NITD008 (50, 100, 250, 500 or 1000 nM) immediately after virus adsorption. Viral titres were quantified at 0 h and 16 h (N=2). Viral titres in NITD008-treated cells at 16 h are shown as individual points with means connected by lines (green, NITD008 P15; black, untreated P22). Mean viral titres of untreated cells at 0 h (input) and of DMSO-treated cells at 16 h are represented by dashed lines.

N=number of independent experiments. Data are shown as individual points, coded by shape according to experimental replicate, with means (connected by lines in F). Statistical tests: one-way ANOVA with Dunnett’s *post-hoc* test (A, D-E), one-way ANOVA with Tukey’s *post-hoc* test (B-C). *, *P* < 0.05; **, *P* < 0.01; ***, *P* < 0.001; ****, P < 0.0001; ns, not significant.
